# The Vagus Nerve conducts viable translocation of gut flora to the lungs that impacts interstitial lung disease severity in mice

**DOI:** 10.64898/2026.05.15.725489

**Authors:** Amit Kumar, Pooja Keerthipati, Humphrey Lotana, Tristan White, Evan Jones, Joseph Prescrille, Tonya Webb, Yuyi Zhu, Andriy Somakhin, Da’Kuawn Johnson, Orest Tsymbalyuk, Marc Simard, Xichun Qin, Yong Ge, Hao Zhang, Shilpa Dilipkumar, Norberto Gonzalez-Juarbe, Wonder Puryear Drake

## Abstract

Communication between gut microbiota and extraintestinal organs is increasingly recognized, yet elucidation of relevant translocation mechanism(s) remains enigmatic. Vagus neuroanatomy and reports of vagal protein transfer to extraintestinal organs suggest that this “superhighway” could translocate bacteria. Here we explore whether the vagus superhighway can translocate bacteria to extraintestinal organs. Gavage of green fluorescent protein–expressing *Escherichia coli* (GFP-*E. coli)* into germ-free (GF) or specific-pathogen free (SPF) C57BL/6 mice yielded high bacillary loads in the stomach and lungs, followed by the heart, stool and peripheral muscles, despite negative blood cultures. Notably, confocal microscopy and culture revealed GFP-*E. coli* within the vagus nerve within five minutes of gavage suggesting rapid translocation. Metagenomic analysis of stool, lung, heart, vagus nerve, and muscle from non-gavaged SPF mice demonstrated significant microbial overlap, supporting that bacterial translocation occurs despite the presence of endogenous microflora. Remarkably, subdiaphragmatic vagotomy performed prior to GFP-*E. coli* gavage resulted in marked reductions of bacterial transduction in the lungs and other extraintestinal organs, except muscle. Furthermore, vagotomy significantly reduced lung fibrosis in SPF mice following intranasal bleomycin administration. In lung cancer patients undergoing lobectomy, vagotomy inhibited postsurgical reductions in forced vital capacity. These findings identify the vagus nerve as a literal gut–lung axis, facilitating viable bacterial translocation and influencing lung severity.

## Introduction

Independent reports support communications between the gut microbial community and remote organs, such as the brain^1^, the lung^2^ and the cardiovascular system^3^. More recent studies corroborate evidence that the gut–lung axis can impact acute and chronic lung diseases. For example, in children, compositionally distinct human neonatal gut microbiota (NGM) are differentially related to relative risk (RR) of childhood atopy and asthma. The highest risk stool composition possessed a distinct fecal metabolome enriched for pro-inflammatory metabolites^4,5^. Similarly in adults, lung or gut dysbiosis, an imbalance of the microbial community towards proinflammatory bacteria, impacts the severity of interstitial lung disease^2,6,7^. Equally noteworthy is evidence that the gut microbiome carries clinical consequences. For example, the maternal gut microbiome impacts the offspring’s risk of allergic disease and asthma, hypothesized to be secondary succinate production by *Prevotella copri* during pregnancy which decreases the offspring’s risk of allergic disease. This in turn promotes bone marrow myelopoiesis of dendritic cell precursors in the fetus^8^. These independent reports relay the importance of gut microbiota to various human diseases.

While the importance of gut microbiota to lung disease severity is not questioned, the mechanism(s) by which gut microbiota or their metabolites get to extraintestinal sites, such as the lung, remain enigmatic. Postulated mechanisms include the following: 1) the microorganisms and microbial-derived products enter the circulation directly^9^ 2) extracellular vesicles, which are secreted from the majority of cell types^10^, facilitate cell–cell communication^11^, 3) intestinal dysbiosis directly damages the physical barrier of the intestinal mucosa by downregulating key tight junction proteins (e.g., Occludin, ZO-1, Claudin-1) triggering “leaky gut”^12^. This allows intestinal toxins (e.g., LPS) and viable bacteria to translocate into the systemic circulation through the mesenteric lymphatic system or portal circulation, worsening systemic inflammation.

The vagus nerve serves as a major bidirectional communication pathway between the brain and internal organs, regulating multiple physiological processes, including autonomic control during states of rest and recovery. Approximately 80% of vagal fibers are afferent, transmitting sensory information from peripheral organs to the brain, while the remaining 20% are efferent fibers that convey motor signals from the brain to visceral tissues^13^. The vagus nerve provides crucial contributions to lung, cardiac and gastrointestinal function. Vagal innervation of the upper respiratory tract was initially described by Galen in the second century AD and further understanding of parasympathetic function was described by Pavlov in the 1800’s^14^. More recent investigation of the vagus nerve influences pathology of various organs: 1) vagal nerve stimulation can inhibit acute lung inflammation through the cholinergic anti-inflammatory pathway^15,16^; 2) vagus nerve stimulation can reduce systemic inflammatory responses to endotoxin^17^ ; 3) vagus nerve stimulation is a promising add-on treatment for treatment-refractory depression, posttraumatic stress disorder, and inflammatory bowel disease^18^. Despite strong clinical association with the vagus nerve and disease pathogenesis, its contribution to translocation of gut flora to various organs remains unexplored.

The emergence of a literal gut-lung axis or the translocation of viable bacteria has not been previously reported. We provide data demonstrating the capacity of the vagus nerve to translocate viable bacteria from the gut to extraintestinal organs, especially the lungs, in mice. In addition, we report that translocation of these viable bacteria to the lungs carries functional and physiologic consequences on interstitial lung disease (ILD) severity.

## Material and Methods

### GFP *Escherichia coli* gavage and quantitative detection by culture

In order to enhance rigor and distinguish gavaged bacteria from commensals, genetically modified *Escherichia coli* containing kanamycin resistance and Green Fluorescent Protein (GFP-*E. coli*), provided as a kind gift from Dr. Tonya Webb (University of Maryland School of Medicine), was used for gavage experiments. Kanamycin resistance is very rare among commensal stool flora of C57Bl/6 mice^19^.

GFP-*E. coli*; 1 × 10⁹ colony forming units [CFU] was administered via gavage into both germ-free (GF) and specific-pathogen free (SPF) C57BL/6 mice. GF and SPF mice who did not undergo gavage served as negative controls. Five minutes after gavage, mice were euthanized, and tissues were collected under sterile conditions for microbial analysis. To minimize contamination, all procedures were performed aseptically. Organs were harvested sequentially according to the predicted lowest bacterial load: blood, thigh muscle, cervical vagus nerve, anterior vagus nerve, aorta, heart, trachea, esophagus, right and left lung lobes, stomach, jejunum, and stool. Tissue homogenates from each of the harvested organs and blood were serially diluted; bacterial culture was performed on Kanamycin supplemented Luria-Bertani (LB) agar for ∼15 h incubation at 37 degrees under aerobic conditions.

Spontaneous resistance to aminoglycosides is uncommon among murine flora^20^. In order to exclude isolation of *E. coli* present as commensal flora, the GFP-*E. coli* strain used to gavage the germ-free and SPF mice carried an extrachromosomal kanamycin resistance gene, as well as green fluorescent protein (GFP).

### Histologic staining and Confocal Microscopy

Sections of the vagus nerve, obtained five minutes after gavage, were fixed in 4% paraformaldehyde (J19943.K2, Thermo Fisher Scientific) and stored at 4°C to localize GFP-positive bacteria within the vagus nerve by confocal microscopy.

### Histologic staining

Antigen retrieval was performed using an antigen retrieval kit (Enzo Life Sciences). Confirmation that the extracted tissue was indeed nerve tissue was performed by staining for neurofilaments. The nerve tissues were then permeabilized with 0.3% Triton X-100 for 30 min, followed by blocking with 10% goat serum for 1 h at room temperature. Samples were then incubated overnight at 4 °C with primary antibody against neurofilament (R&D Systems). After washing, tissues were incubated with fluorescently conjugated secondary antibody (Alexa Fluor 647 goat anti-rat; R&D Systems). Finally, samples were mounted on glass slides using antifade mounting medium containing DAPI (Vector Laboratories). Images were acquired with the Abberior Facility Line microscope at the Confocal Microscopy Core at UMB. Confocal imaging was performed using an Olympus 60x/1.42 STED-UPLXAPO60xO oil immersion lens. The images for gavaged and non-gavaged samples were acquired using the same acquisition parameters and images from both conditions shown in the figure are display-adjusted similarly to reduce the tissue background in the green channel and image analysis was performed using Fiji software (version 2.17.0).

### DNA extraction and metagenomic analysis by organ systems

Fecal pellets, aorta, muscle, right upper lung, right lower lung, and vagus nerve samples were collected from SPF mice in each housing cohort. Genomic DNA (gDNA) was extracted using the DNeasy Blood & Tissue Kit (Qiagen, Valencia, CA) according to the manufacturer’s protocol. DNA concentration and integrity were assessed using the Bioanalyzer 2100 system (Agilent, Santa Clara, CA). Genomic DNA libraries were constructed for sequencing on the Illumina platform using the NEB Ultra Express FS DNA Library Prep Kit (New England Biolabs, Ipswitch,MA). Libraries were sequenced on an Illumina NovaSeq 6000 using 150bp PE reads targeting 50M reads per sample (Illumina, San Diego, CA).

Paired end sequences were trimmed, quality-filtered, and host contaminants were removed in KneadData (v.0.7.6) using default values and the C57BL mouse reference database. Shotgun metagenome sequencing of bulk amplified genome DNA was performed. Quality control of each metagenome was performed using tools from the BBMap software package (http://sourceforge.net/projects/bbmap/) and the bioBakery3 suite^21^. Identical duplicates were removed using the Clumpify tool in “dedupe” mode allowing 0 substitutions. BBDuK was run to remove reads from the PhiX spike-in and synthetic molecules (k=31). Kneaddata (v0.10.0) was run to trim low-quality bases, remove adapters, discard short reads, and filter human reads (--trimmomatic-options=" ILLUMINACLIP:<adapters.fa>:2:30:10 SLIDINGWINDOW:4:20 MINLEN:60" -db hg37dec_v0.1). Taxonomic profiling was performed using tools from the Kraken suite^22^. Kraken was run with default parameters using the Standard database. Bracken was executed with read length of 150bp. Taxa outside of the domains Archaea and Bacteria were excluded from downstream analyses. Compositions of sample types were examined for relative abundances at the phylum. A 0.01% minimum abundance filter was applied to remove low abundance species prior to visualization. Sequences of tissue microbiota have been deposited into BioProject ID PRJNA1466953.

#### Bioinformatics

Metagenomic sequencing reads were assembled using metaSPAdes v3.10^23^. Following assembly, scaffold headers were standardized to include sample-specific prefixes using SeqKit^24^ and a simplified annotation format (SAF) file was subsequently generated to facilitate downstream feature quantification. To estimate scaffold coverage and abundance, the original paired-end reads were mapped back to the final assembly using BoSPFie2^25^. An index was constructed using boSPFie2-build, and alignment was performed in "very-sensitive" mode with end-to-end alignment parameters to maximize sensitivity. Post-alignment processing was conducted using SAMtools^26^. Resulting SAM files were converted to BAM format. Alignments were strictly filtered to retain only properly paired alignments. An adaptive selection strategy was applied to the filtered BAM files to retain contigs meeting robust detection thresholds. Read abundance was quantified using featureCounts^27^ in paired-end mode. Coding sequences were predicted by Prodigal^28^. Taxonomic classification was performed using the CAT (Contig Annotation Tool) package^29^ with default parameters. The resulting abundance tables were processed using a custom Python 3 workflow utilizing the Pandas library. Taxonomic identifiers were normalized and data aggregated by species identity, summing abundance counts across all sample columns. To reduce noise, the dataset was filtered to exclude species with a cumulative abundance count of less than 10.0 across all samples. Plots and statistical analyses were made with GraphPad Prism (Version 11.0.0) and R Studio (Version 2026.01.1+403).

### Vagotomy followed by bacterial quantification

To confirm the importance of the vagus nerve to extraintestinal bacterial translocation, GFP-*E.coli* gavage, culture and quantification was conducted as described above after subdiaphragmatic vagotomy of the anterior vagus nerve (AVN) or sham operation (SHAM). Mice were anesthetized with xylazine (14mg/kg) and ketamine (140mg/kg). Buprenorphine E.R. (0.075mg/kg) was injected intraperitoneal as a presurgical analgesics. Mice were placed in the supine position, and the abdomen area was shaved and cleaned with alcohol and betadine solution. Following skin disinfection, abdominal muscles were incised 1 cm along the ventral median line to expose the liver. The liver was then carefully moved upwards to expose the esophagus and stomach. We transected the left trunk of the vagus nerve subdiaphragmatic portion under a surgical microscope and sewed the abdominal muscles and skin and administered analgesic carprofen (4mg/kg) as previously described^30^. The same procedure was performed in sham-operated mice, except no vagus nerve transection occurred. Three weeks after vagotomy, mice were gavaged with GFP-*E. coli* for quantification of organ bacillary load post vagotomy or sham operation.

### Vagotomy followed by bleomycin induction of murine lung fibrosis

Vagotomy was performed as described above. Bleomycin induction of pulmonary fibrosis, Ashcroft histologic scoring and lung collagen quantification were conducted as previously described^2^. Mice were housed in groups of three per cage, and all mice in each cage received the same treatment. Randomization was not used to allocate experimental units to control and treatment groups. Determination of sample size was based upon a minimum of five mice per group. On average, there were 14 mice per group. All murine procedures were performed according to the protocol approved by the Institutional Animal Care and Use Committee at University of Maryland School of Medicine (protocol #AUP00000179; ABSL-2 facilities). No criteria were set for excluding animals from analysis a posteriori. We excluded any wild-type mice from the immune analysis and Sircol assay who died before 14 days, which is the timeframe required to develop lung fibrosis. Lungs were harvested for histology, flow cytometry, or single cell isolation as reported^2^. The outcome measures were histologic scoring and collagen quantification.

### Clinical investigation of the role of vagotomy on forced vital capacity

A single-center, randomized controlled trial of lung cancer patients, aged equal to or greater than 65 years of age, undergoing elective video-assisted thoracoscopic lobectomy and lymphadenectomy were randomized at a 1:1 ratio to undergo a sham procedure (control group [n=16]) or transection of the pulmonary branches of the vagus nerve that innervate the bronchial stump plus the caudal-most large pulmonary branch of the vagus nerve (n=16)^31^. This clinical trial followed the principles of the Declaration of Helsinki and Consolidated Standards of Reporting Trials (CONSORT) Guidelines. The protocol was approved by the Ethics Committee of Authors’ hospital. Informed consent was obtained prior to enrollment from all participants (Approval No. XYFY2019-KL179-01). The trial is registered at Clinicaltrials.gov (NCT04247997). While the original trial reported significant reductions in chronic cough in the population receiving vagotomy, the impact on forced vital capacity (FVC) between the sham operated and vagotomized patients was not reported. Here, we report the impact of vagotomy or sham-operation on the FVC change in cancer patients 65 years old and above, undergoing lobectomy.

## Statistical Analysis

Data analysis and graph generation were performed using GraphPad Prism version 11.0.1 (GraphPad Software). Results are expressed as mean ± standard error (SE). Statistical significance between two groups was evaluated using an unpaired Student’s t-test. For comparisons across multiple groups, a one-way analysis of variance (ANOVA) followed by Tukey’s multiple comparisons test was applied. Each data point represents an individual mouse. A P value < 0.05 was considered statistically significant. All murine experiments were conducted using cohorts of 14 mice, depending on sample availability. Lung, aorta, muscle, vagus nerve and stool microbiota analyses were performed once in a cohort of five mice.

## Results

Rapid dissemination of GFP-*E. coli* to lungs and other extraintestinal sites following gavage of germ-free mice.

Known as the wandering nerve, the vagus nerve is the longest cranial nerve in the body with the broadest territorial innervation. Through the dorsal motor nucleus of the vagus nerve, there are efferent and afferent branches throughout the body including the esophagus, the heart, the lungs, the stomach, the small and large intestines, the gallbladder, the liver, the pancreas, the kidneys, the spleen, the thymus, and the adrenal gland^32,33^. The vagus nerve is a major conduit lung through the afferent nerves, P2ry1 and Npy2r^34^. The vagus nerve has been shown to translocate proteins, such as alpha-synuclein, from the stomach to the brain^35^. We hypothesized that bacteria may be translocated to extraintestinal sites as well. We assessed for a literal gut-lung axis following gavage of GFP-*E. coli* from the stomach to extraintestinal sites.

Remarkably, within five minutes of gavage of 1x10^9^ GFP-*E. coli*, bacteria were cultured from multiple extraintestinal sites, such as the lung, heart, vagus nerve, stomach, peripheral thigh muscle and stool of gavaged GF mice (n=13). Jejunal involvement of bacteria was also noted. Bacteria were absent from the corresponding organs of all non-gavaged mice (n=14), resulting in significant distinctions in bacteria isolation among gavaged and nongavaged GF mice (p<0.0001, Welch’s t test) (Figures 1A-G; Supplemental Figure 1). Surprisingly, the blood was culture negative for GFP-*E. coli* in both gavaged and non-gavaged mice, suggesting that hematogenous transmission from the gut to the lung and other extraintestinal sites was unlikely (Figure 1H). Quantitatively, approximately 80% of the gavaged bacteria were present in the stomach and jejunum (data not shown). Notably, the lungs represented the most common extraintestinal site for bacterial translocation, followed by the heart, vagus nerve and thigh muscle (Figure 1I-K). Notably, there were no significant quantitative differences among individual members of an organ system, such as the right middle lobe, right upper lobe and Left lung lobe, or heart versus aorta (Figure 1J; Supplemental Figure 1).

**Figure 1.**
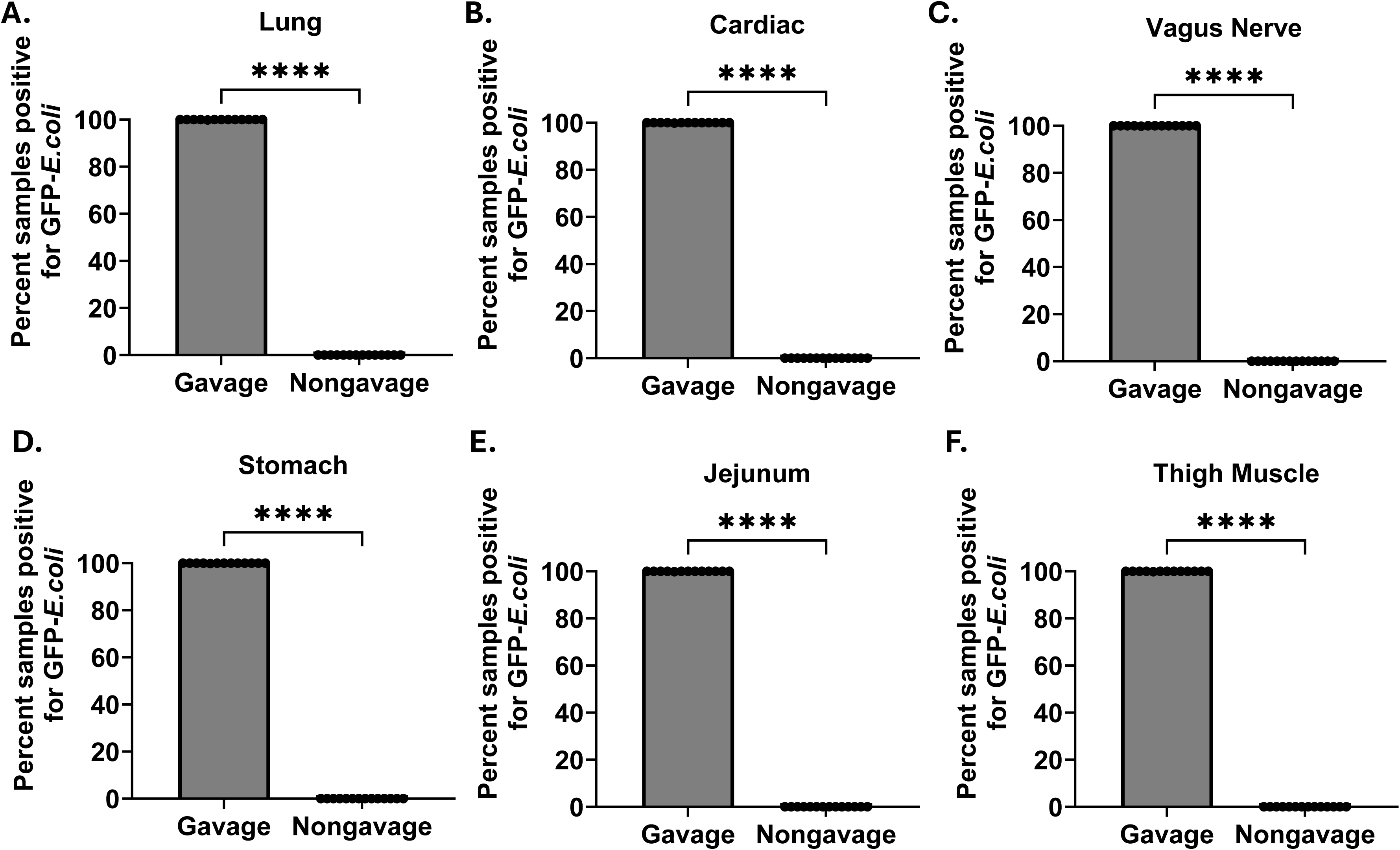
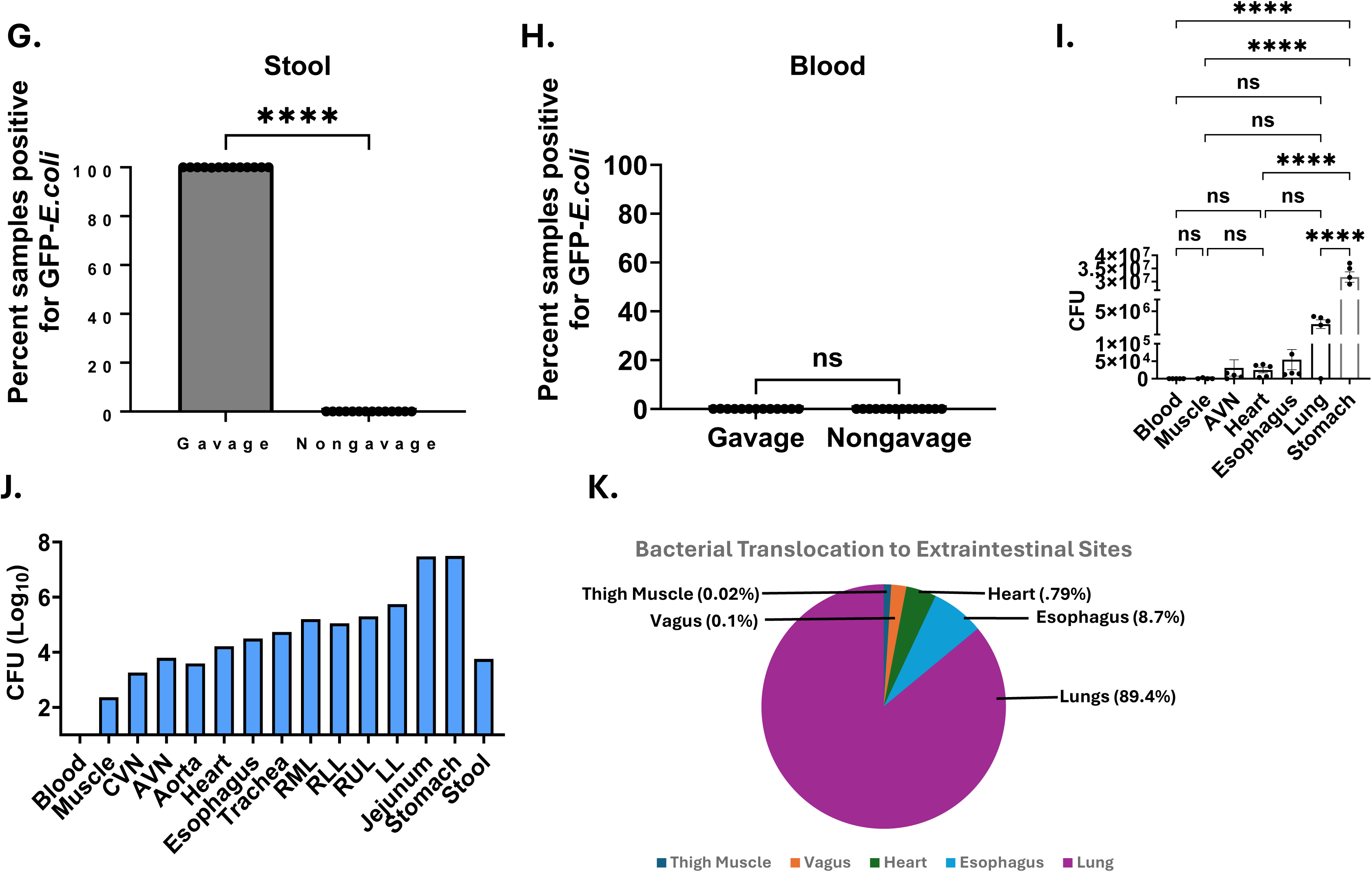
Rapid dissemination from the gut to extraintestinal sites following orally-administered GFP-*E. coli* in germ-free mice. Germ-free mice were gavaged (100μl) with GFP-*E. coli* (1 × 10⁹ CFU) or PBS (sham). Mice (n = 5–14) were euthanized 5 minutes post-gavage, and tissues were harvested and cultured for bacteria on plates containing kanamycin. Qualitative and quantitative analysis for bacteria was performed ∼15 hours after organ harvest. The presence of CFU was compared between gavaged and non gavaged miced mice for each organ: **A)** Lung, **(B)** Cardiac, **C)** Vagus nerve, **D)** Stomach, **E)** Jejunum, **F)** Thigh Muscle, **G)** Stool and **H)** Blood. **I)** Colony-forming unit (CFU) counts recovered from the indicated organs (n=organs of 5 mice per group). **J)** CFU counts in blood, thigh muscle, carotid vagus nerve (CVN), anterior vagus nerve (AVN), aorta, heart, esophagus, trachea, right middle lobe (RML), right lower lobe (RLL), right upper lobe (RUL), left lung (LL), jejunum, stomach and stool, demonstrating the distribution of viable bacteria across tissue compartments. **K)** Pie chart summarizing the proportion of bacterial transfer originating from the gut, illustrating the relative contribution of gut-derived microbes to extraintestinal colonization. Statistical significance was assessed using unpaired t-test for gavaged and non-gavaged organ involvement, as well as one-way ANOVA with Tukey’s multiple comparisons. ****P < 0.0001, and; ns, not significant.

### Rapid dissemination of GFP-*E. coli* to lungs and other extraintestinal sites following gavage of mice with commensal flora

In order to assess if the presence of commensal flora impacted bacterial extraintestinal translocation, we repeated the same extraintestinal qualitative and quantitative bacterial assessment five minutes following gavage of GFP-*E. coli* in C57Bl/6 SPF mice. Qualitatively, bacteria were present in extraintestinal sites, such as lung, heart, vagus nerve, thigh muscle, and stool of gavaged mice but absent from all non-gavaged mice (p<0.0001, Welch’s t test) (Figure 2A-G). Again, no GFP-*E. coli* were detected in the blood of gavaged or non-gavaged mice (Figure 2H). Quantitative analysis revealed significant distinctions among bacteria present in the organs of gavaged SPF mice (Figure 2I-J). The highest number of bacteria in the extraintestinal sites were again detected in the lungs, followed by esophagus, heart, vagus nerve, stool and thigh muscle (Figure 2K).

**Figure 2.**
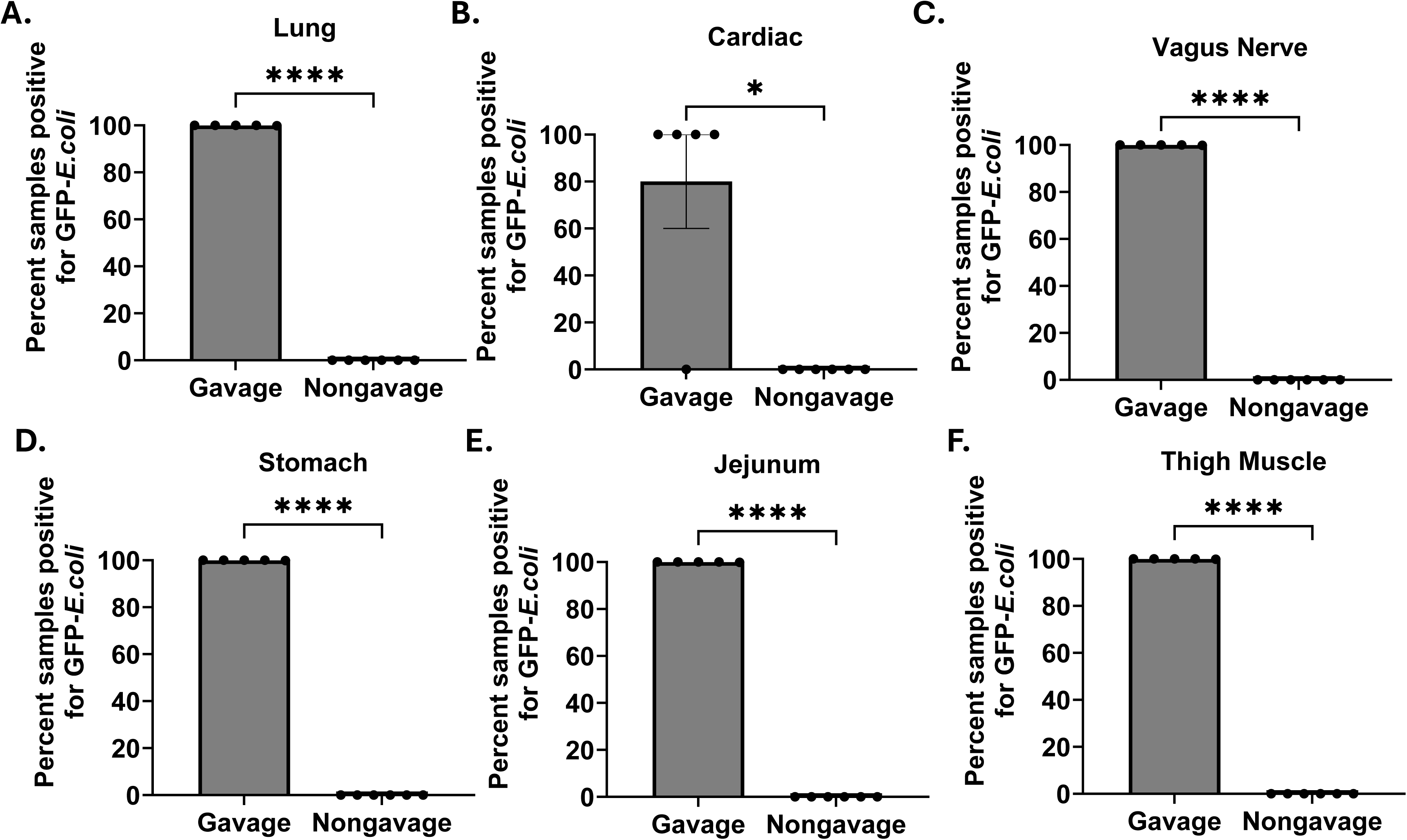
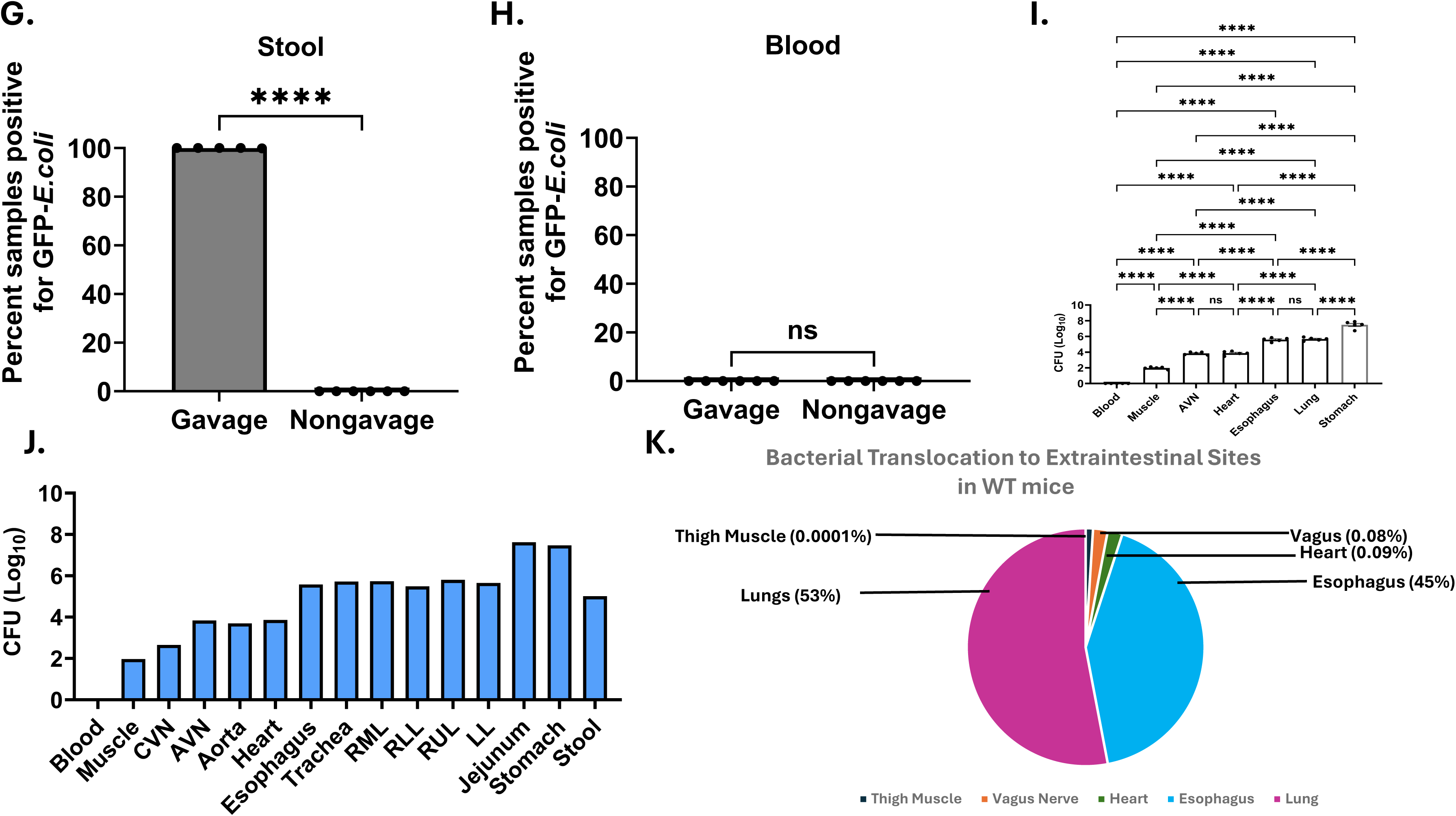
Rapid extraintestinal dissemination of orally administered GFP *E. coli* in SPF mice. SPF mice were gavaged (100μl) with GFP *E. coli* (1 × 10⁹ CFU) or PBS (sham). Mice (n = 5–6) were euthanized 5 minutes post-gavage, and tissues were harvested and cultured for bacteria on plates containing kanamycin. Qualitative and quantitative analysis for bacteria was performed ∼15 hours after organ harvest. The presence of CFU was compared between gavaged and non gavaged miced mice for each organ system: **A)** Lung, **B)** Cardiac, **C)** Vagus nerve, **D)** Stomach, **E)** Jejunum, **F)** Thigh Muscle, **G)** Stool and **H)** Blood. **I)** Colony-forming unit (CFU) counts recovered from the indicated organs (n=5 each group). **J)** CFU counts in blood, thigh muscle, carotid vagus nerve (CVN), anterior vagus nerve (AVN), aorta, heart, esophagus, trachea, right middle lobe (RML), right lower lobe (RLL), right upper lobe (RUL), left lung (LL), jejunum, stomach and stool, demonstrating the distribution of viable bacteria across tissue compartments. **K)** Pie chart summarizing the proportion of bacterial transfer originating from the gut, illustrating the relative contribution of gut-derived microbes to extraintestinal colonization. Statistical significance was assessed using unpaired t-test for gavaged and non-gavaged organ involvement, as well as one-way ANOVA with Tukey’s multiple comparisons. ****P < 0.0001, and; ns, not significant.

Akin to the observation in GF mice, there were no significant differences between individual tissues within an organ system; the left lungs had similar quantity as the right lobes, as were the aorta and the heart (Figure 2J; Supplemental Figure 2). Also, akin to the observation in GF mice, the lungs were the most common site of extraintestinal translocation (Figure 2K). We also assessed for bacterial translocation among gastric extraintestinal organs and noted significant translocation to the kidney, liver, spleen and pancreas in gavaged SPF mice (Supplemental Figure 3).

### Confocal microscopy visualization of GFP-*E. coli* within the vagus nerve

We began by performing confocal microscopy to assess for the presence of GFP-*E. coli* within the vagus nerve five minutes after gavage. We tested vagus serve sections from gavaged (n=5) and non-gavaged (n=6) mice. GFP signals were present in the neuronal cytoplasm of the vagus nerve of gavaged mice but absent from all non gavaged mice (Figure 3A). We also cultured the vagus nerve on LB plates containing kanamycin and again noted GFP-*E. coli* from the vagus nerve of gavaged mice but no bacteria were isolated from non-gavaged mice (Figure 3B-C). Anatomically, the vagus nerve is comprised of ∼80% afferent neuronal fibers (sensory transmission from peripheral sites to central nervous system) and ∼20% efferent neuronal fibers (transmission from CNS to peripheral motor neurons). We hypothesize that there would be quantitative distinctions along the vagus nerve. Sections of the vagus nerve were obtained from the anterior vagus nerve (AVN) and the cervical vagus nerve (CVN). The AVN is the portion of the vagus nerve that enters through the diaphragm then splays along the less curvature of the stomach; the CVN is the section of the vagus nerve that is located with the carotid sheath, posterior to the carotid artery. The AVN had a significantly higher average bacterial concentration than the CVN (p=0.01, Welch’s t test) (Figure 3D). Remarkably, these findings indicate rapid translocation of viable bacteria, likely along the afferent fibers of vagus nerve, with bacterial concentrations highest at sites closer to the gavage site.

**Figure 3.**
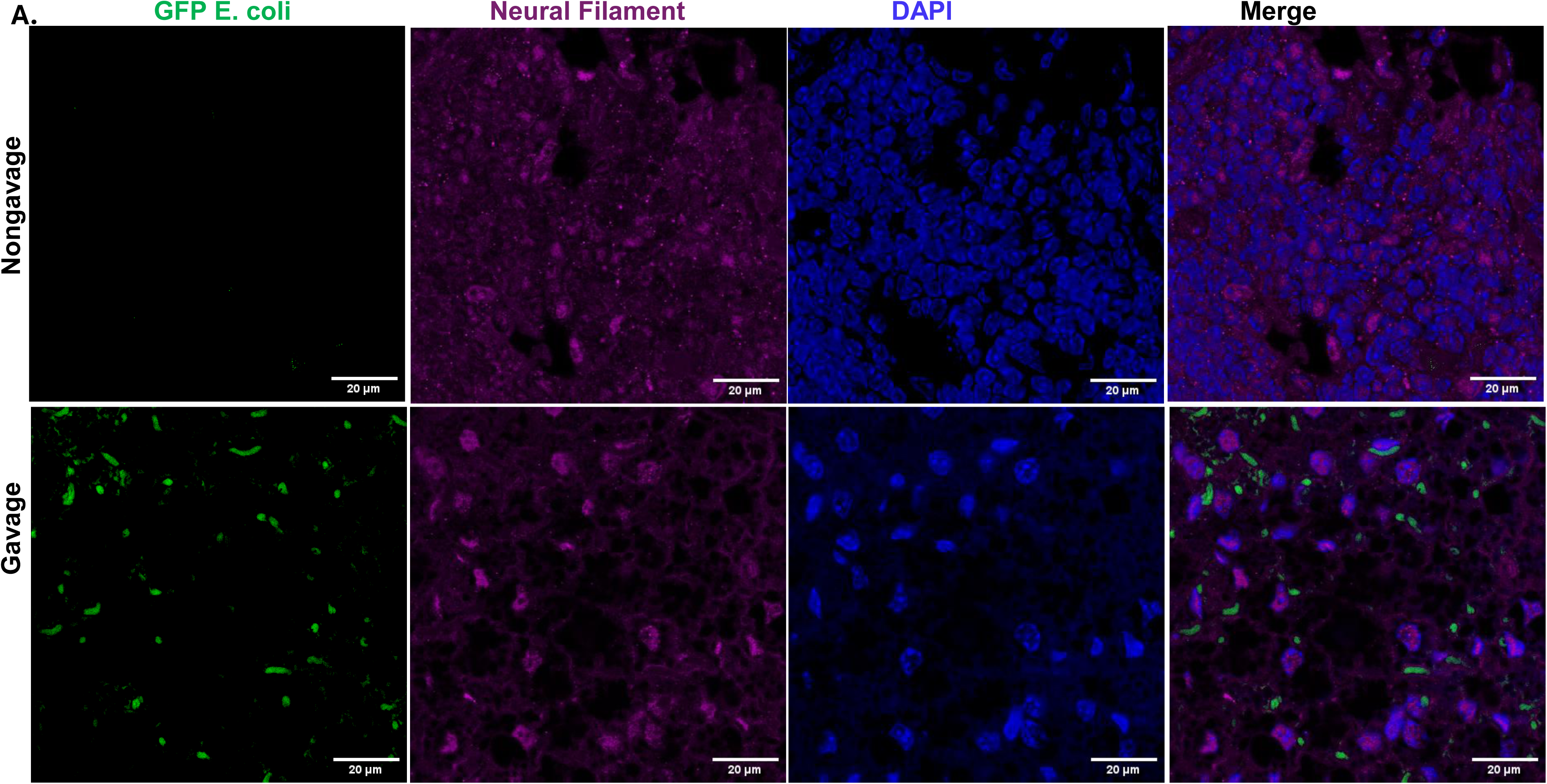
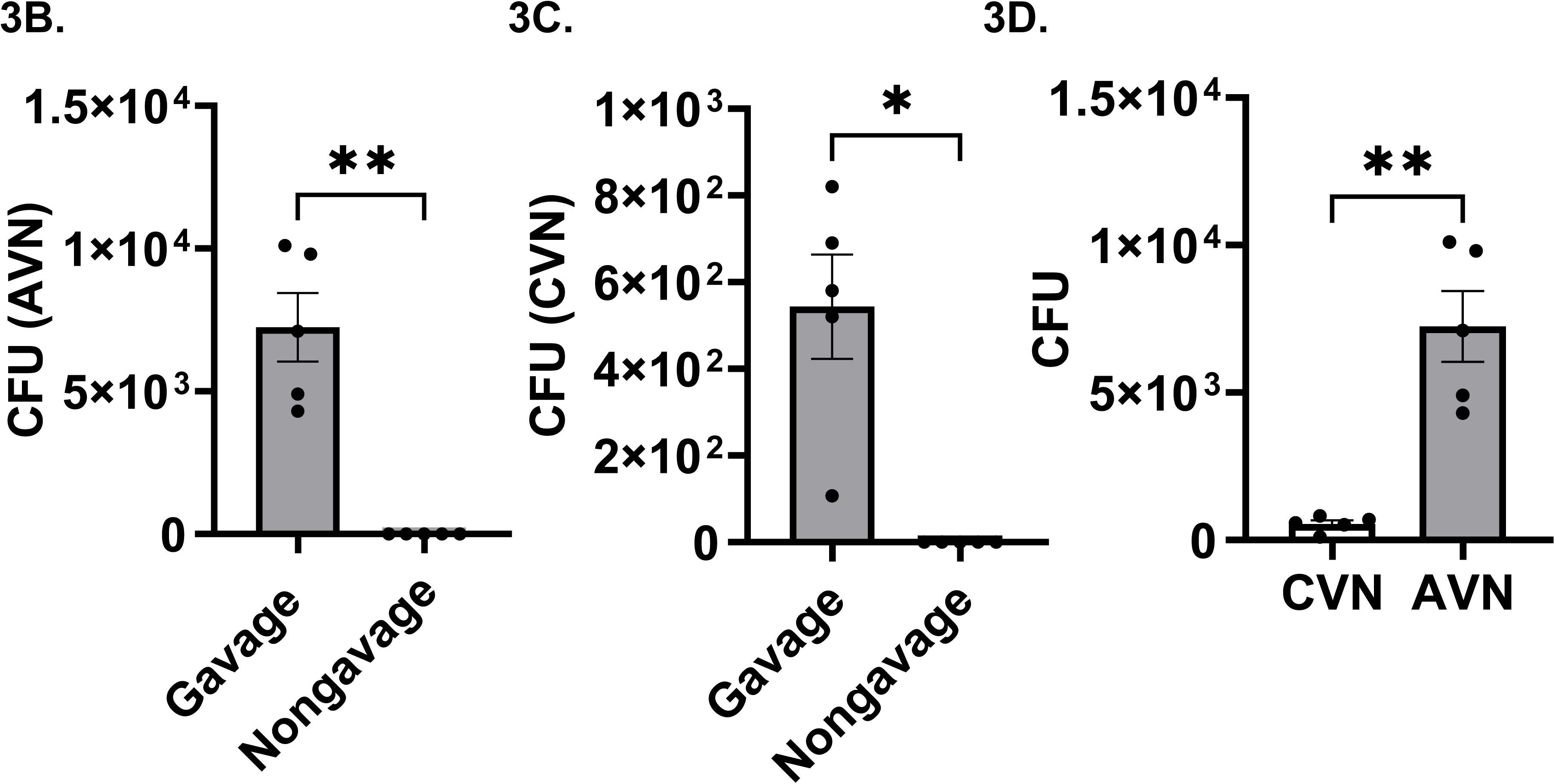
Confocal microscopy and culture of vagus nerve for GFP-*E. coli*. **A)** Confocal microscopy of the vagus nerve 5 minutes after gavage with GFP-*E. coli* in germ-free mice shows bacterial localization within neuronal cytoplasm. Nerve fibers were stained with neurofilament (cyan), and nuclei were counterstained with DAPI (blue). Scale bar, 20 μm. Quantification of colony-forming units (CFU) in the gavage (n=6) and non gavaged (n=5) mice **B)** Anterior vagus nerve (AVN), **C)** Cervical vagus nerve (CVN) and **D)** CFU difference in gavage CVN and AVN, 5 minutes after gavage with GFP-*E. coli*. Data are presented as mean ± SEM, with each symbol representing an individual mouse. Statistical significance was determined using an unpaired *t*-test. *P < 0.05 and **p<0.01.

### Metagenomic analysis of lung, heart, vagus nerve, thigh muscle and stool

The studies thus far support active translocation of bacteria to extraintestinal sites following gavage. To assess if this occurs spontaneously, we conducted metagenomic analysis to assess of intestinal and extraintestinal sites to assess for congruence among the microbial communities.

### Shared microbial signatures across tissues reflect possible translocation of organisms from a common source

Here we aimed to determine the compositional differences between the microbiome present in the stool, vagus nerve, aorta, muscle and lung tissues. Tissues from SPF mice housed in ABSL2 conditions were carefully excised and processed for metagenomic sequencing with control non-tissue samples added to determine background contamination. Assessment of bacterial species richness and relative abundance by Shannon and Chao indexes showed the stool had the highest richness and abundance with the aorta, right upper lung (RUL), right lower lung (RLL), muscle, and vagus showing similar profiles (Figure 4A-C). Then we used a principal coordinates analysis (PCoA) to determine and visualize similarities or differences between the tissue and stool samples. PERMANOVA confirmed that microbial community composition differed significantly between groups (R² = 0.888, *p* = 0.001), being the stool the only group compositionally different from the rest. No significant differences in dispersion were observed, indicating that these differences reflect true compositional variation rather than heterogeneity in variance (Figure 4D). Subsequently, we tested the overall relative abundance of the top 20 most abundant taxa (> 1% abundance) across all samples. We observed that microbial communities were dominated by *Curtobacterium flaccumfaciens*, which accounted for a vast amount of the relative abundance. Contributions from additional taxa, including *Akkermansia muciniphila*, *Exiguobacterium sp*, and *Staphylococcus aureus* were observed at lower abundances (Figure 4E). A heatmap of log-transformed abundances demonstrated consistent presence and abundance of non-*Curtobacterium flaccumfaciens* taxa compared to the stool (Figure 4F). Analysis of taxonomic overlap revealed extensive sharing of microbial taxa across all samples, including vagus nerve, lung (RUL and RLL), muscle, aorta, and stool. UpSet analysis^36^ demonstrated that the largest set of taxa belonged to the stool sample (matching its higher diversity), however, there was an observed overlap of 14 taxa that were present in all tissues supporting the notion of a shared core microbiome (Figure 4G-H). A relative abundance plot of the 14 shared organisms showed the high abundance of two taxa (*Curtobacterium flaccumfaciens* and *Akkermansia muciniphila*) and the lower abundance of the other 12 shared organism (Figure 4H). *Curtobacterium* spp are plant bacteria rarely shown to be pathogenic in animals, which was likely present due to the large number of plants present in the lab. We reanalyzed the data following the removal of *Curtobacterium flaccumfaciens* and noted the same consistency of community flora across the various organs, many of which were known enteric flora such as *Clostridium*, *Bacteroides* and *Klebsiella* spp (Supplemental Figure 4A, B). Taken together, the shared microbial signatures across tissues reflect possible translocation of organisms from a common source, the gut.

**Figure 4:**
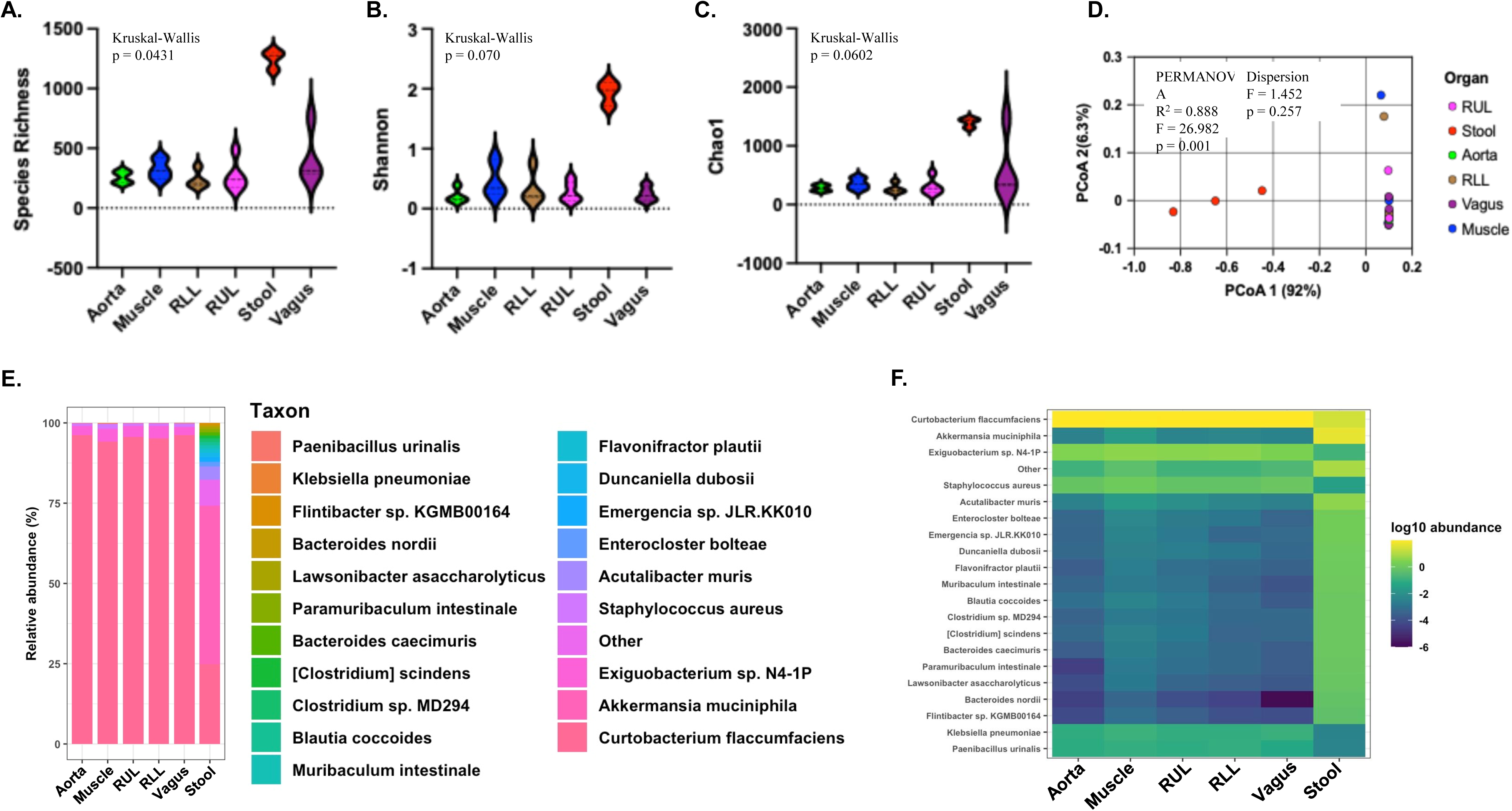
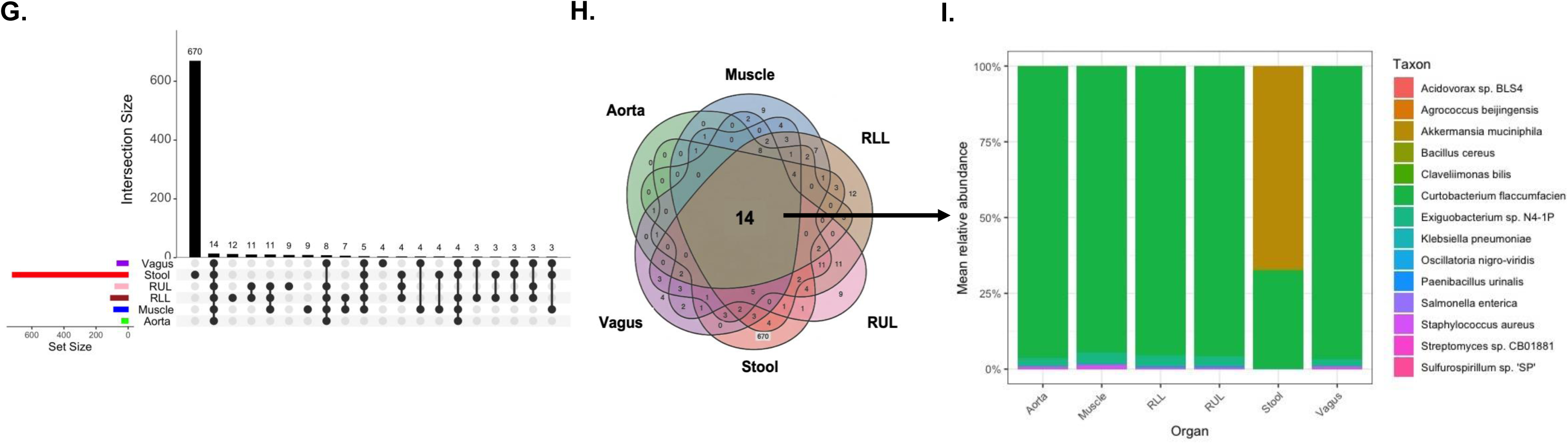
Vagus nerve has a core microbiome indistinguishable from muscle, aorta and lung tissues. A–C) Alpha diversity across vagus nerve, stool, right upper lung (RUL), right lower lung (RLL), muscle and aorta samples, shown using species richness, Shannon and Chao analyses. **D)** Principal coordinates analysis (PCoA) based on β-diversity distances demonstrating separation of samples by tissue type; PERMANOVA indicated significant differences between groups (R² = 0.888, F = 26.982, p = 0.001), with no significant differences in dispersion (p = 0.257). **E)** Relative taxonomic composition of top 20 most abundant species shown as stacked bar plots. **F)** Heatmap of top 20 log10-transformed taxon abundances across samples. **G)** UpSet plot and **H)** Venn diagram with heatmap summarizing shared and unique taxa across tissues, alongside mean relative abundance across tissue types.

### Vagotomy reduces bacterial translocation to extraintestinal sites

To examine the necessity of vagal integrity to the rapid, extraintestinal distribution of gavaged GFP-*E. coli*, we performed subdiaphragmatic vagotomy or sham surgery (opening, then immediate closure of the abdominal wall without vagotomy) in SPF mice. The left AVN was surgically removed and the right AVN remained intact. Following surgical healing (∼21 days later), both murine cohorts were gavaged with GFP-*E. coli*; tissues were harvested five minutes post-gavage for quantitative analysis of GFP-*E. coli* CFU following culture on kanamycin-supplemented LB plates.

Remarkably, the organs harvested from vagotomized mice displayed marked reductions in bacterial loads from extraintestinal sites compared to sham-operated controls, including the lungs, heart, cervical vagus nerve, and stool (Figure 5A-J). Reductions in the subsections of each organ, such as the RLL, RML and left lobe of the lung revealed similar significant reductions (Figure 5A-J). There were some notable exceptions to post-vagotomy reductions in bacillary load; for example, no significant reductions in the peripheral thigh muscle was noted (Figure 5K). The bacillary loads within the stomach and jejunum were not impacted by vagotomy (Figure 5L-M); however, significant reductions in the stool were noted following vagotomy (Figure 5J). This suggests that the gavaged microorganisms reach the stool via the vagus nerve and not simply gastric propulsion. Consistent with our previous findings, no CFU were detected in the blood of either group (Fig. 5N). Taken together, these data support that the vagus nerve plays a critical role in facilitating the rapid distribution of gut flora to extraintestinal sites.

**Figure 5.**
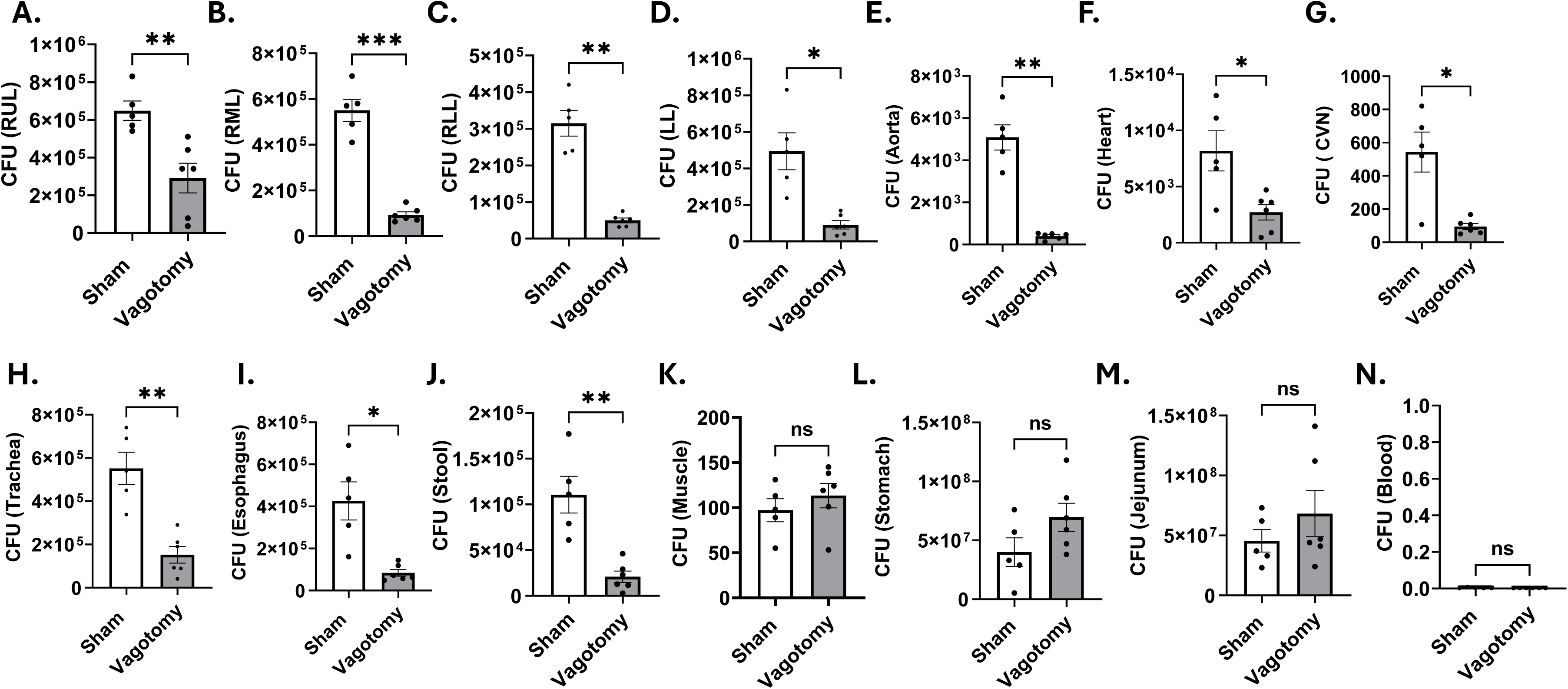
Vagotomy limits the rapid extraintestinal dissemination of orally delivered GFP-*E. coli*. SPF mice received subdiaphragmatic vagotomy (n = 6) or sham surgery (n = 5). three week later, mice were gavaged with 1 × 10⁹ CFU GFP-*E. coli*. Five minutes after gavage, tissues were harvested and plated on kanamycin-supplemented LB agar; bacterial burdens were determined after 15 h. Quantification was performed in muscle **A)** Right upper lobe (RUL); **B)** Right middle lung (RML); **C)** Right lower lobe (RLL); **D)** Left lung (LL); **E)** Aorta; **F)** Heart; **G)** Carotid vagus nerve (CVN); **H)** Trachea; **I)** Esophagus; **J)** Stool; **K)** Muscle; **L)** Stomach; **M)** Jejunum and **N)** Blood. Data are shown as mean ± SE, with each symbol representing one mouse. Statistical significance was determined using an unpaired *t*-test. *P < 0.05; **P < 0.01; ***P < 0.001; ns, not significant.

### Disruption of vagus nerve translocation reduces bleomycin-induced lung severity

Prior reports implicate dysbiotic gut flora in interstitial lung disease (ILD) severity^2,7^. To determine the importance of vagal bacterial translocation to ILD severity, we induced lung fibrosis through intranasal bleomycin inoculation to vagotomized or sham-operated SPF mice housed in ABSL-2 conditions. Control mice receiving intranasal saline served as a negative control. While blinded to the surgical assignment, histologic Ashcroft scoring and quantitative assessments of lung collagen content via the Sircol assay were conducted in the sham-operated and vagotomized mice.

Ashcroft scoring of hematoxylin and eosin (H&E) and trichrome staining of right middle lobe lung sections revealed significantly less fibrosis in vagotomized mice compared to sham-operated mice (Figure 6A, B). Remarkably, significant reductions in lung collagen content was also noted in the vagotomized mice, compared to the sham-operated (Figure 6C; p=0.01, One way Anova with Tukey’s). The lung collagen content of vagotomized mice was similar to the mice receiving normal saline, and significantly less than bleomycin-treated sham-operated or mice without surgical manipulation. The sham-operated mice had the same elevated lung collagen content as mice without surgical manipulation (Figure 6C).

**Figure 6.**
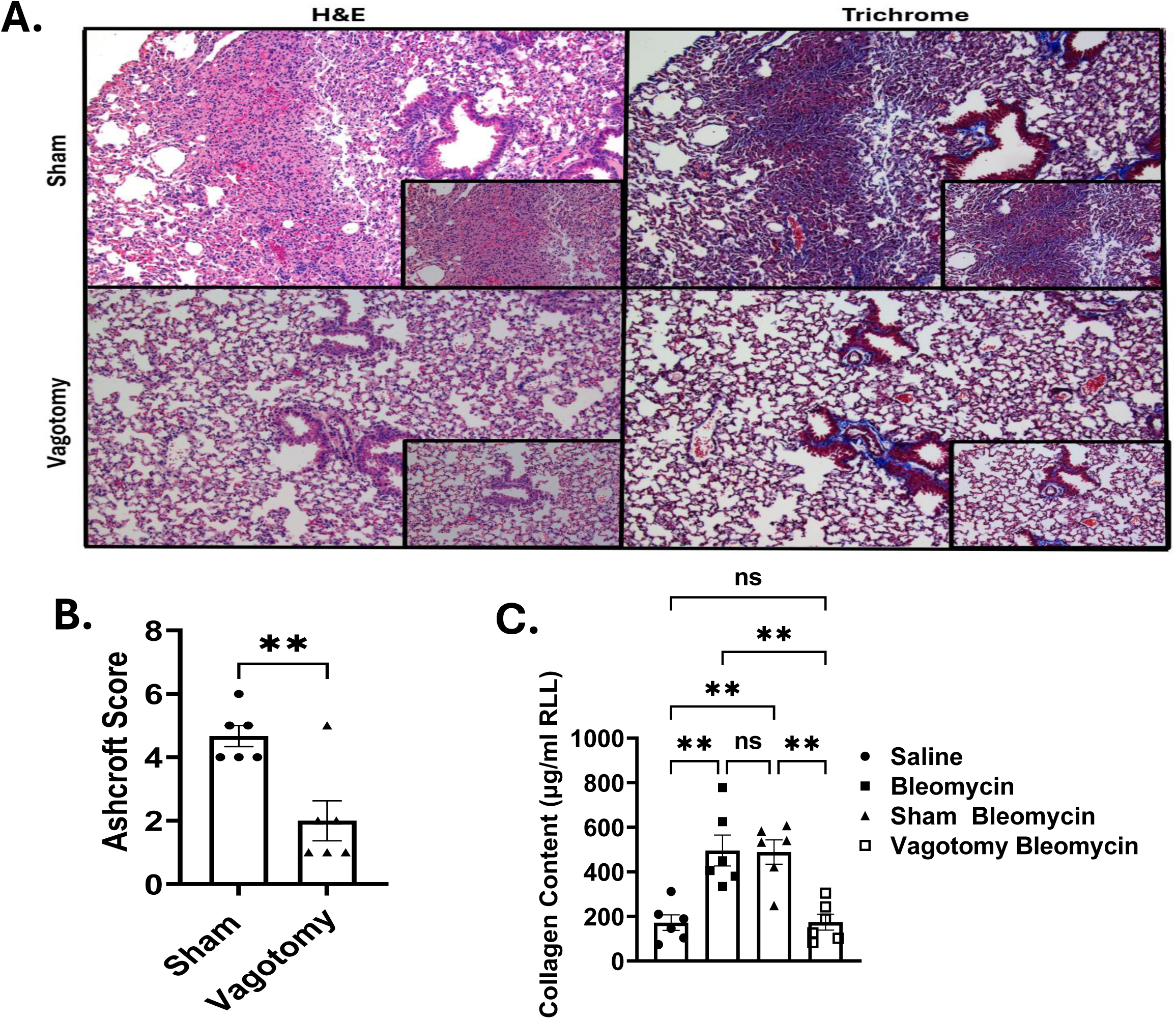
Vagotomized SPF mice are protected from lung fibrosis. Lung collagen content in SPF, SPF-vagotomized, and sham-operated mice housed in ABSL-2 facilities and intranasally inoculated with 30 μl containing 0.04 U bleomycin. Control animals received 30 μl saline intranasally. **A)** Representative images for H&E and trichrome-stained lungs of Sham and Vagotomy mice, 14 days after bleomycin injury with Scale bar, 100 μm. **B)** Ashcroft Scoring of H and E-stained lungs by housing conditions following bleomycin administration to mice (n = 6 per group). **C)** Soluble collagen levels in the right lower lobe (RLL) were quantified using the Sircol assay. Data are presented as mean ± SE, with each point representing an individual mouse (n = 24; 6 per group). Statistical significance was assessed with multiple cohorts using a one-way ANOVA with Tukey’s multiple comparisons and Ashcroft scoring using Welch’s t-test. **P < 0.01, and P < 0.01; ns, not significant.

Independent reports relay the importance of the gut-lung axis in lung cancer outcomes^4,37^, including the impact of gut microbiota on lung function in lung cancer patients^38^. We wanted to assess for physiologic relevance of vagotomy in humans. The vagus nerve is part of the cough reflex, which is common after lobectomy for lung cancer. In a single-center, randomized controlled trial, adult patients >65 years of age, undergoing elective video-assisted thoracoscopic lobectomy and lymphadenectomy were randomized at a 1:1 ratio to undergo a sham procedure (control group) or transection of the pulmonary branches of the vagus nerve that innervate the bronchial stump plus the caudal-most large pulmonary branch of the vagus nerve. Transection of the pulmonary branches of vagus nerve prevented chronic cough following lobectomy. Lung function tests were also obtained. We investigated the impact of vagus nerve transection on forced vital capacity (FVC) among patients undergoing sham operation (n=16), compared to those who had a transection of the pulmonary branch of the vagus nerve (n=16). We noted a significant FVC decline among those who experienced lobectomy with a sham vagotomy (p=0.02, Welch’s t test) (Figure 7A). However, the cohort of patients undergoing lobectomy with vagotomy had no significant FVC decline (p=0.08, Welch’s t test) (Figure 7B). These findings suggest that vagotomy may protect against FVC decline among older cancer patients undergoing lobectomy.

**Figure 7.**
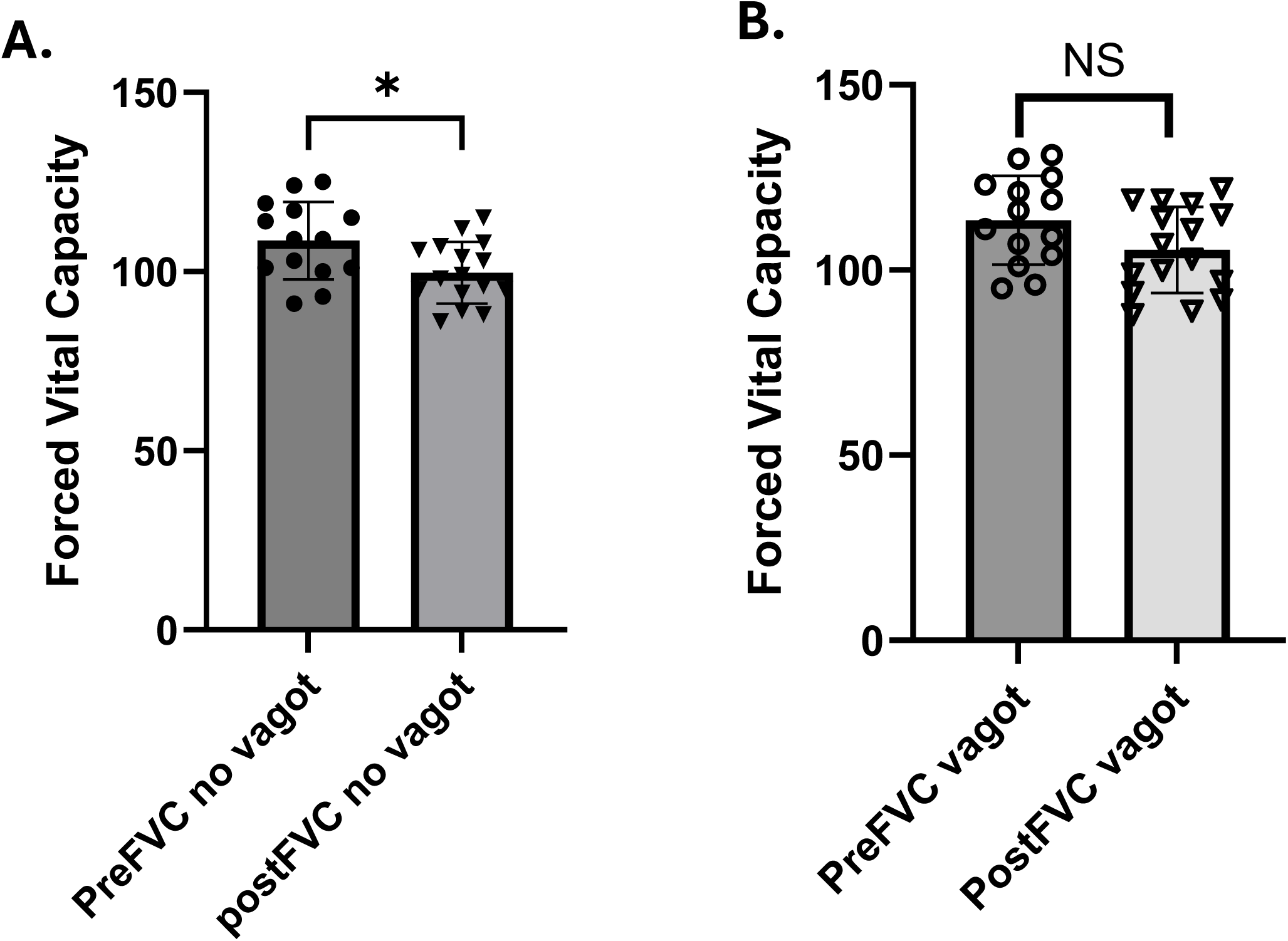
Vagotomy protects against significant decline in FVC among lung cancer patients undergoing lobectomy. All available baseline and post-operative forced vital capacity (FVC) values were obtained among 32 patients undergoing lobectomy for lung cancer. **A)** Patient who did not undergo vagotomy had a significant FVC decline (n=16) (p=0.02, Welch’s t test), **B)** whereas a nonsignificant decline was noted in vagotomized patients (n=16) (p=0.08. Welch’s t test).

**Table 1.**
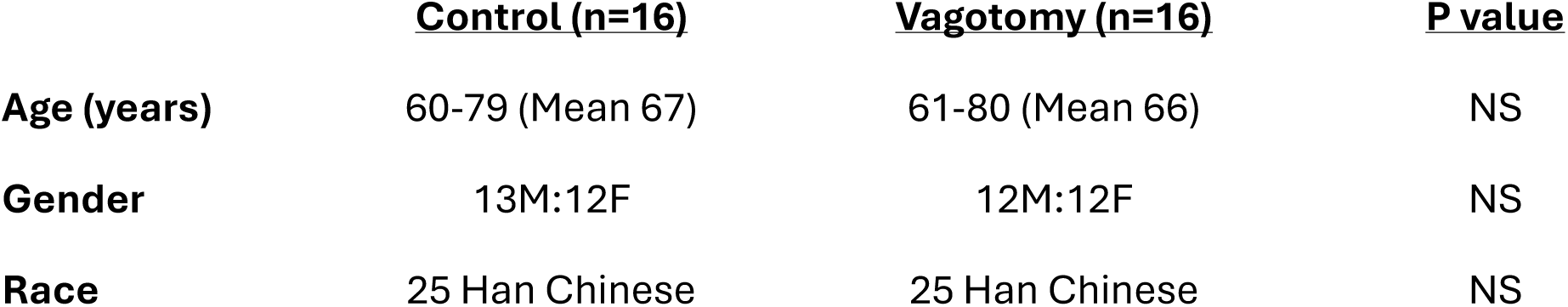
Demographic Data of Patients undergoing Lobectomy with or without vagotomy.

|  | <u>Control (n=16)</u> | <u>Vagotomy (n=16)</u> | <u>P value</u> |
| --- | --- | --- | --- |
| **Age (years)** | 60-79 (Mean 67) | 61-80 (Mean 66) | NS |
| **Gender** | 13M:12F | 12M:12F | NS |
| **Race** | 25 Han Chinese | 25 Han Chinese | NS |

These murine and human findings support that an intact vagus nerve carries a functional significance on lung severity; the murine studies support that the vagus nerve serves as a literal gut–lung axis (Figure 8).

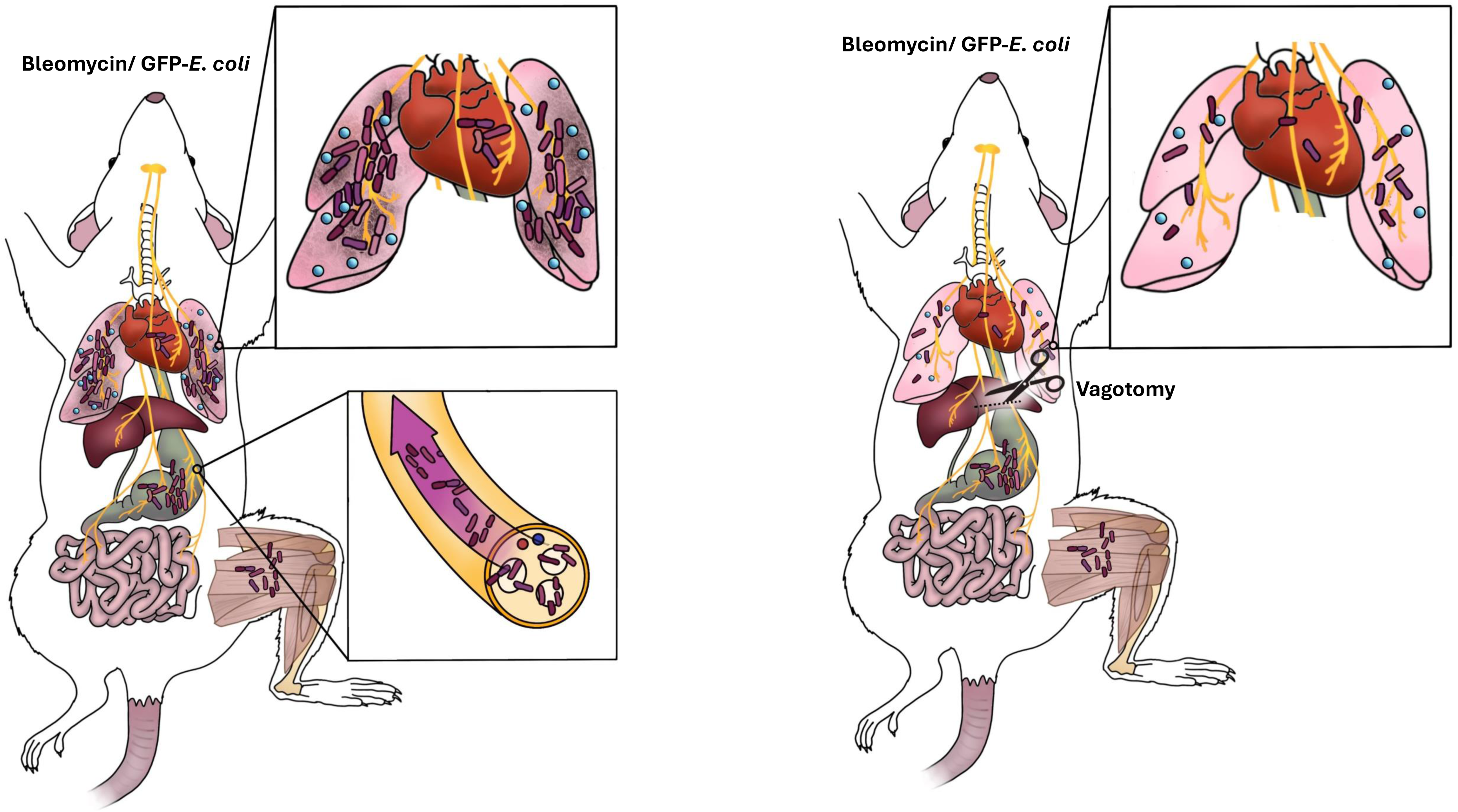

## Discussion

These studies provide in vivo confirmation of the capacity of the vagus nerve to rapidly translocate viable bacteria to extraintestinal sites, of which the lung is the most common. These findings demonstrate that the vagus nerve functions as a literal gut-lung axis and, most notably, that the transport of viable microorganisms can exacerbate interstitial lung disease severity.

The translocation of viable organisms to extraintestinal sites is rapid, occurring as early as 5 minutes after gavage. *GFP-E. coli* replicates every 20 minutes. In order to minimize organ CFU to bacillary replication, we chose time points for organ harvest less than of the time necessary for a single round of bacterial replication. It is interesting that in the lungs were the most common site of extraintestinal translocation in both the GF and SPF mice (Figure 1, 2). We also observed translocation to the heart, which may explain the connection of dysbiotic gut flora to cardiac conditions such as atrial fibrillation^39,40^. Equally noteworthy is the consistent observation that when GFP-*E. coli* gavage was performed at lower concentrations, such as 10^3^ and 10^6^; no evidence of extraintestinal bacterial translocation was noted (data not shown), suggesting that a quantitative threshold must be present in order for bacteria to translocate to extraintestinal organs.

Vagotomy significantly reduced bacterial flora in each organ except peripheral thigh muscle, the blood (which was sterile by culture analysis), stomach and small intestine (Figure 5). For the stomach and small intestine, translocation likely does not occur as the stomach was inoculated and the jejunum likely was inoculated through propulsion. However, bacillary loads were reduced in the stool, suggesting that the vagal nerve contributes to stool bacillary load in the terminal gastrointestinal tract. Consideration of rapid translocation to the lung and stool suggest that use of those sites to clear waste products may be one benefit of vagal translocation. The metagenomic studies revealed very similar pathogenic and nonpathogenic microbial patterns in the lungs, vagus nerve and muscle suggest that the vagus nerve does not selectively translocate bacteria. Bacterial extracellular vesicles are key mediators involved in bacteria-to-host communication^41^. Future investigations to assess their role in vagal translocation is warranted.

Vagotomy supported the hypothesis that the vagus nerve’s capacity to translocate bacteria carries functional consequences, such as worsening ILD severity. Vagotomy abrogated its capacity to do so by reducing the translocation of bacteria (Figure 5, 6). Other investigations suggest similar importance for the vagus nerve to lung diseases, such as lung cancer^4,37^ and asthma^42,43^. It has been reported that inoculation of healthy mice with gut microbiota from mice undergoing unpredictable chronic stress activates the vagus nerve and induces early and sustained changes in both serotonin and dopamine neurotransmission pathways in the brainstem and adult hippocampus^44^. We only observed gut translocation of bacteria to the brain in one mouse, suggesting that at the bacterial concentration used to gavage mice, vagal translocation to the brain is rare. We also did not observe reductions in peripheral muscles following vagotomy. There are reports of transneuronal transport of herpes simplex 1 can occur from spinal nerves innervating the muscle to the central nervous system^45^. It may be the vagus nerve:intersynapse: spinal nerves may be responsible for translocation to muscles and vagotomy alone is not sufficient to inhibit translocation.

Vagotomy is not as common a surgical practice as it was in the 1940’s when it was the gold standard for treatment of peptic ulcer disease^46–48^. Further studies are warranted to understand how vagal translocation of flora impacts lung disease. If vagotomy is selected as a mechanism, careful consideration of the site is important, as well as confirmation that the patient does indeed have dysbiotic gut flora. Chemical vagotomyto reduce bacterial translocation should be a possibility. Also, understanding of how the vagus nerve serves as a superhighway with physiologic consequences, ie could it just as easily translocate bacterial flora producing short chain fatty acids to reduce ILD severity. While these findings are provocative, there are limitations to this investigation. For example, while we investigated more than 12 extraintestinal sites, we did not assess every extraintestinal site. There may be other areas that have not been identified, such as the skin. It is possible that our brain assessment could have been more thorough; we assessed in the regions of the brain near the midbrain (region of insertion of the vagus nerve) and noted only two bacterial CFU in one murine brain (data not shown). Gut-brain translocation of bacteria has been described but the work has not been published following peer review^49^. Another limitation is that we did not investigate for viral or fungal “translocation”. It is likely that viral translocation does occur, as translocation of HSV 1 from peripheral muscle to the CNS has been reported^45^. Third, this investigation focused on translocation from the gut to extraintestional sites. The vagus nerve has bidirectional transmission^50,51^. Assessing for transmission from the lung to the GI tract would also enhance understanding and could possibly explain why 1/3 of patients with pneumonia have diarrhea^52,53^.

These findings carry striking implication for the transport of gut flora to sites outside of the gastrointestinal track. They open doors to numerous downstream investigations, including determining the relevance of this pathway to enteric pathogens inducing gram-negative pneumonia, age-related atrial fibrillation^39,40,54,55^ and other diseases impacted by organ fibrosis, such as cirrhosis and chronic kidney disease. These findings support new therapeutic interventions for diseases impacted by gut dysbiosis. Investigations to assess the vagus nerve as a literal gut-heart axis, gut-liver axis and gut-kidney axis are warranted.

## Acknowledgements

This work was supported by grants R01 HL157533, NIH K24 HL127301-01 and Ellen Dreiling Research Fund Endowment to W.P.D, R01AI061061 to T.W., R01 AI168313 to N.G.J, R37NS141786 to M.S. The germ-free mice are bred and housed at the University of Louisville Core facility supported by NIH/NIGMS CoBRE grant (P20GM125504). The SPF mice were obtained from the University of Maryland School of Medicine Animal Facility. We thank the University of Maryland School of Medicine Institute for Genome Sciences for assistance with the metagenomic processing. We also thank the University of Maryland CIBR Confocal Microscopy Core for assistance with confocal imaging.

## Author contributions

A.K and P.K. designed and performed experiments, analyzed data, and wrote the manuscript; H.L., T.W., E. J., T.W., Y.Z., A.S., D. J., O.T., performed experiments; M. S. provided intellectual input and edited the manuscript; X.Q., Y.H., HZ collected the clinical data; N.G.J. performed the metagenomic analysis and wrote the manuscript. performed experiments. W.D. generated the hypothesis, contributed to the intellectual content and writing/editing of the manuscript.

## Competing interests

The authors declare no competing interests.

## Additional information

### Supplementary information

The online version contains supplementary material.

**Corresponding author:** Correspondence and requests for materials should be addressed to Wonder P. Drake.

---

**Supplemental Figure 1 . Extraintestinal dissemination of orally administered GFP *E. coli* in germ free (GF) mice.** The presence of CFU was compared between gavaged (n=13)and nongavaged (n=14) mice using unpaired t-test for each organ: (A), RUL (Right upper lung) (B), RML (Right middle lung) (C), RLL (Right lower lobe) (D), LL (Left lung) (E), CVN (Carotid vagus nerve) (F), AVN (Anterior vagus nerve) (G), Heart (H), Aorta (I), Trachea and (J) Esophagus. Statistical significance was determined using an unpaired *t*-test. ****P < 0.0001.

**Supplemental Figure 2 . Extraintestinal dissemination of orally administered GFP *E. coli* in SPF mice.** The presence of CFU was compared between gavaged (n=5)and nongavaged (n=6) SPF mice using unpaired t-test for each organ: (A), RUL (Right upper lung) (B), RML (Right middle lung) (C), RLL (Right lower lobe) (D), LL (Left lung) (E), CVN (Carotid vagus nerve) (F), AVN (Anterior vagus nerve) (G), Heart (H), Aorta (I), Trachea and (J) Esophagus. Statistical significance was determined using an unpaired *t*-test. *P < 0.05; ****P < 0.0001.

**Supplemental Figure 3 . Extraintestinal dissemination of orally administered GFP *E. coli* in SPF mice.** The presence of CFU was compared between gavaged and non gavaged SPF mice using t-test for each organ: (A), Kidney (B), Liver (C), Spleen (D), Pancreas (n=10 each group). Statistical significance was determined using an unpaired *t*-test. *P < 0.05; ***P < 0.001; ****P < 0.0001.

**Suppplemental Figure 4. Relative taxonomic composition of top 20 most abundant species shown as stacked bar plots.** Composition of top flora detected in the lungs, heart, vagus, peripheral thigh muscle with (A) and without (B) inclusion of stool flora, after removal of the soil bacterium, *Curtobacterium flaccumfacien*.

## Notes

### Competing Interest Statement

The authors have declared no competing interest.

