## Supplementary material for "The Vagus Nerve conducts viable translocation of gut flora to the lungs that impacts interstitial lung disease severity in mice": Supplemmental figure

Supplemental Figure 1

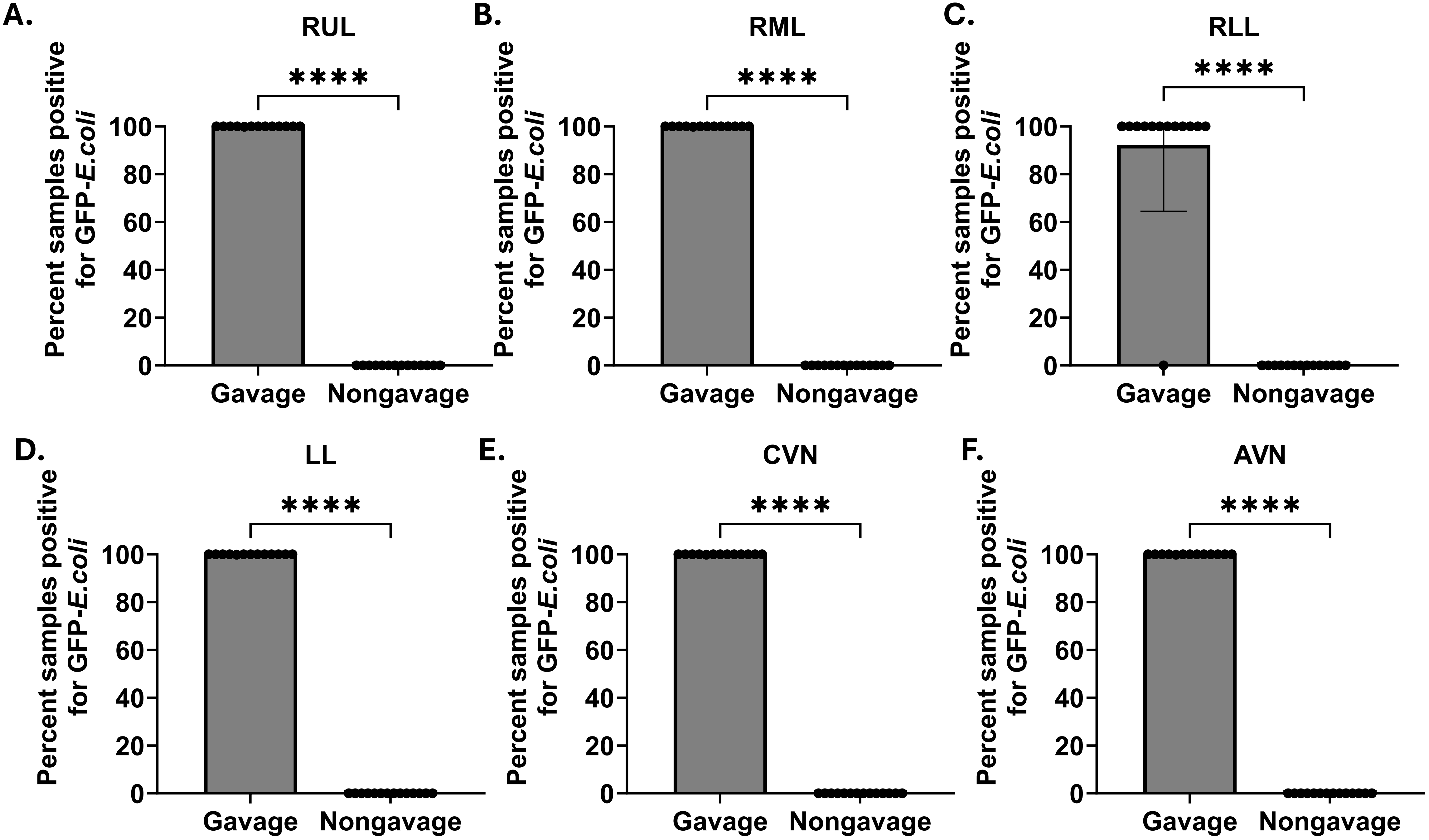

Supplemental Figure 1

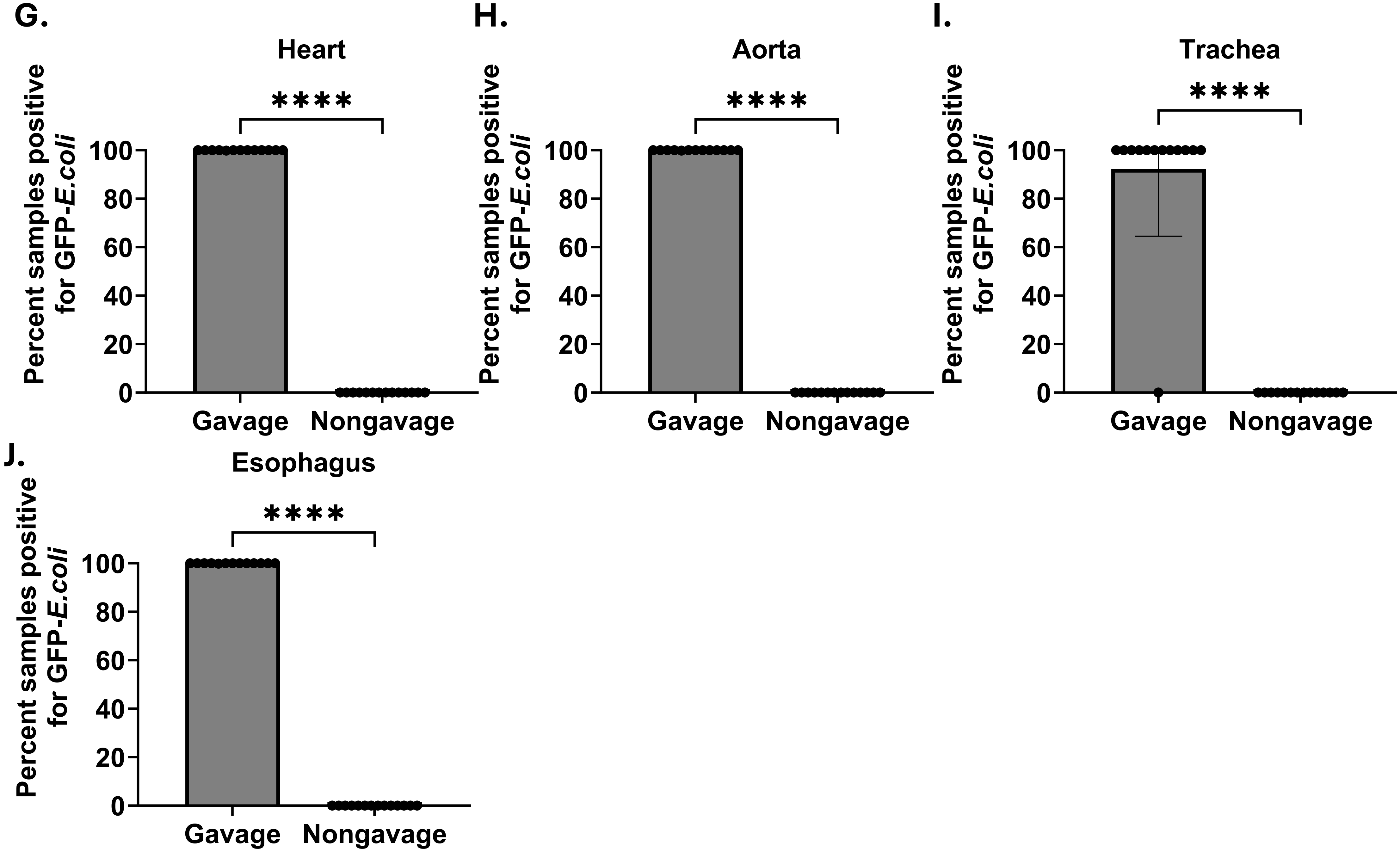

Supplemental Figure 2

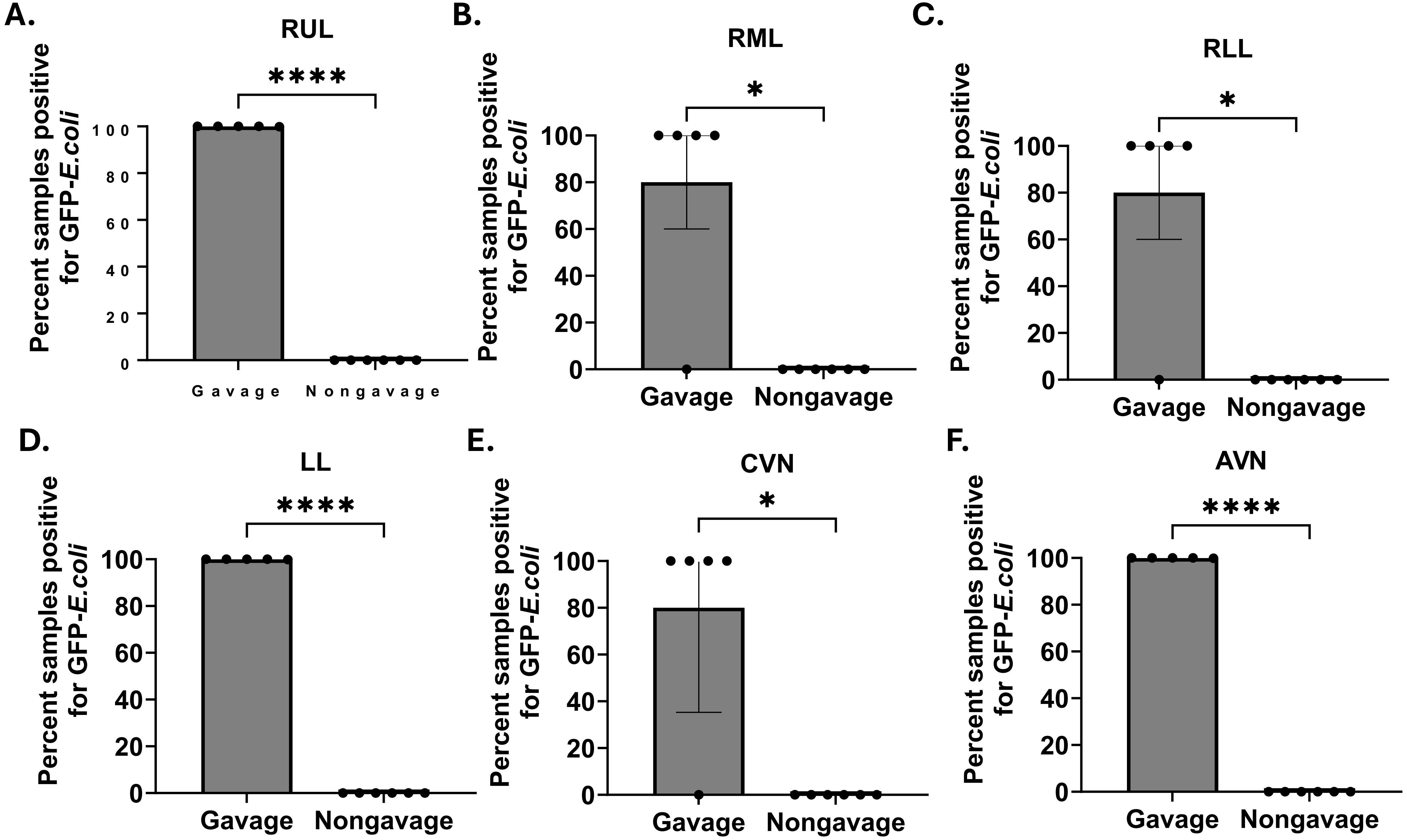

Supplemental Figure 2

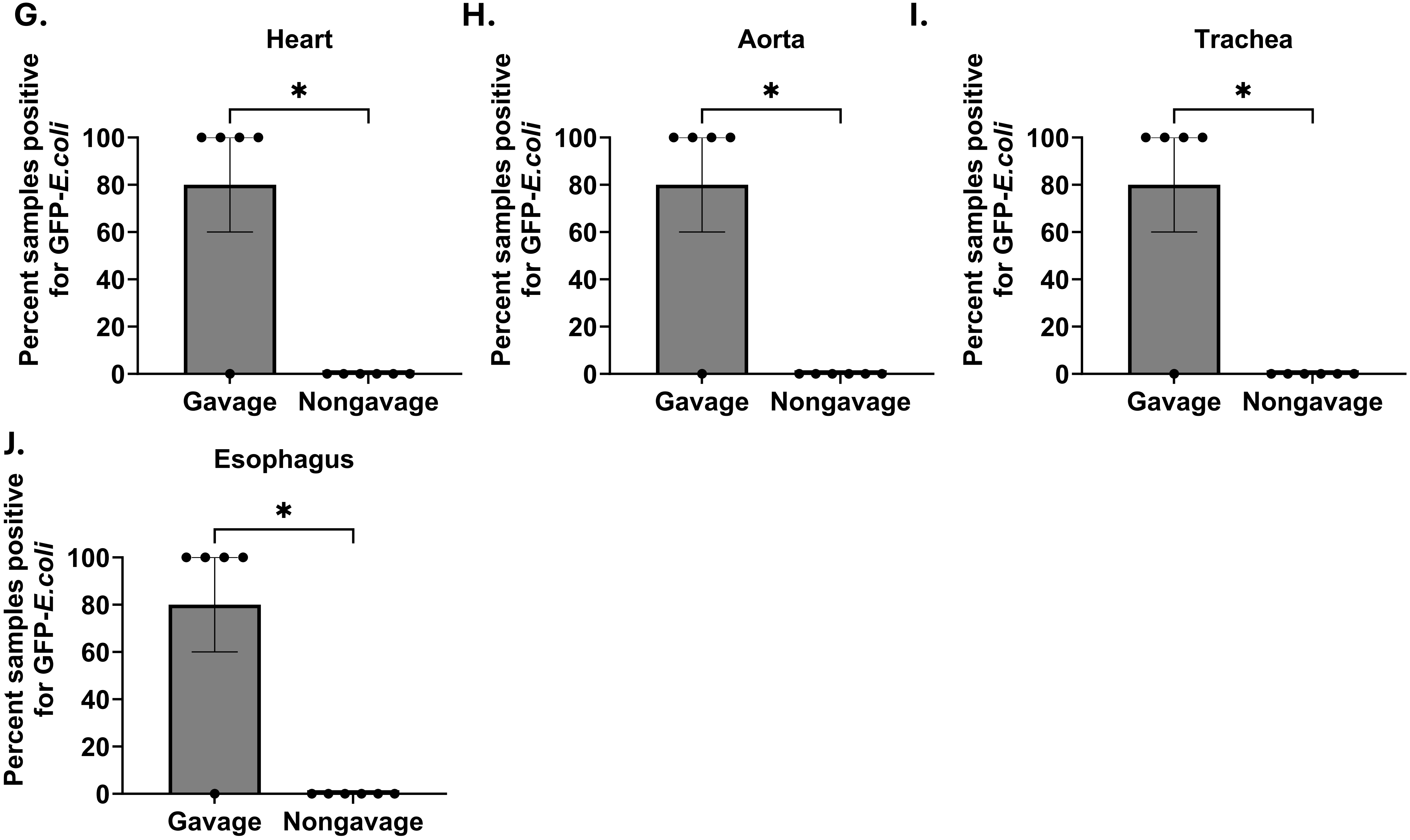

Supplemental Figure 3

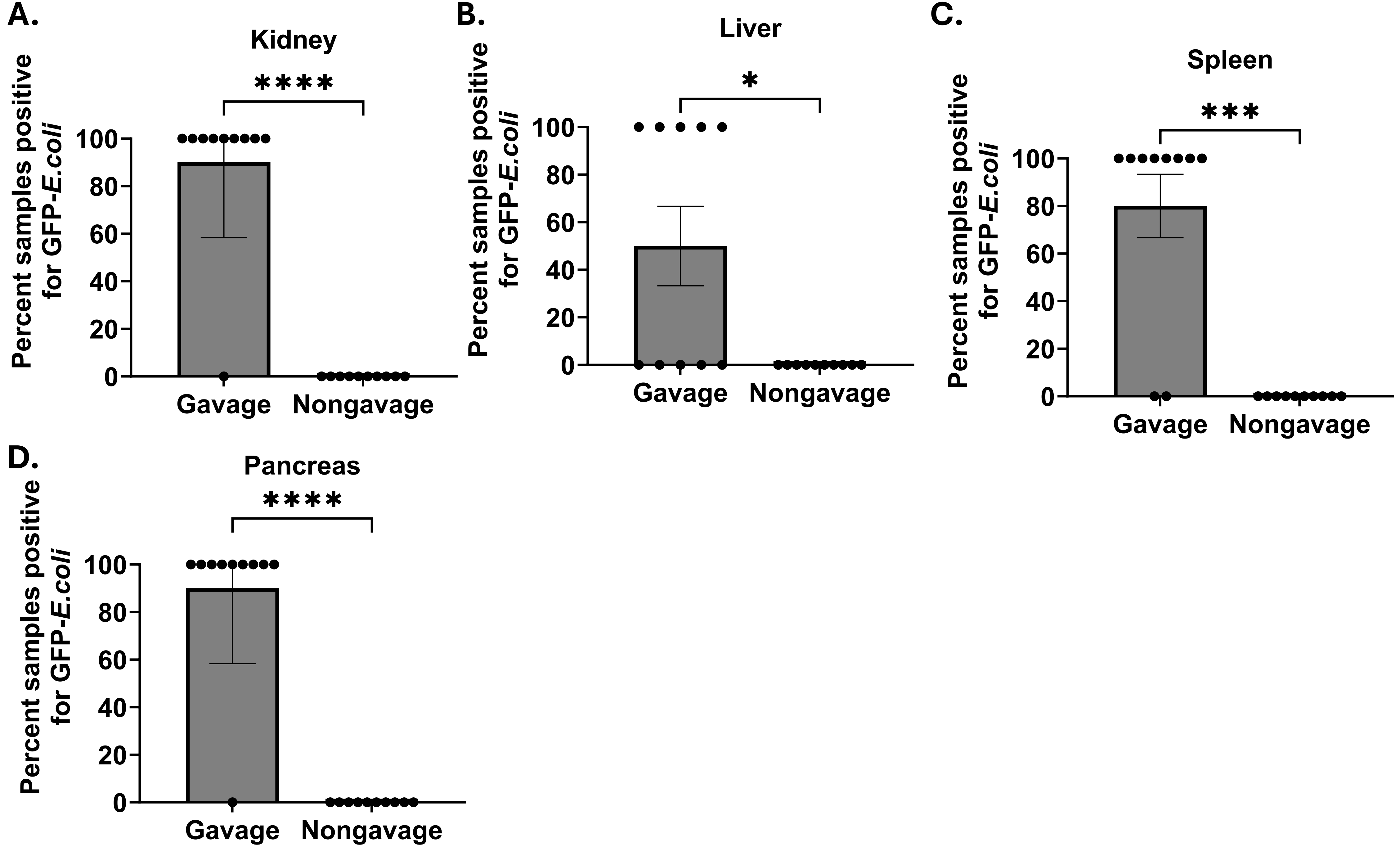

### Supplemental Figure 4

A.

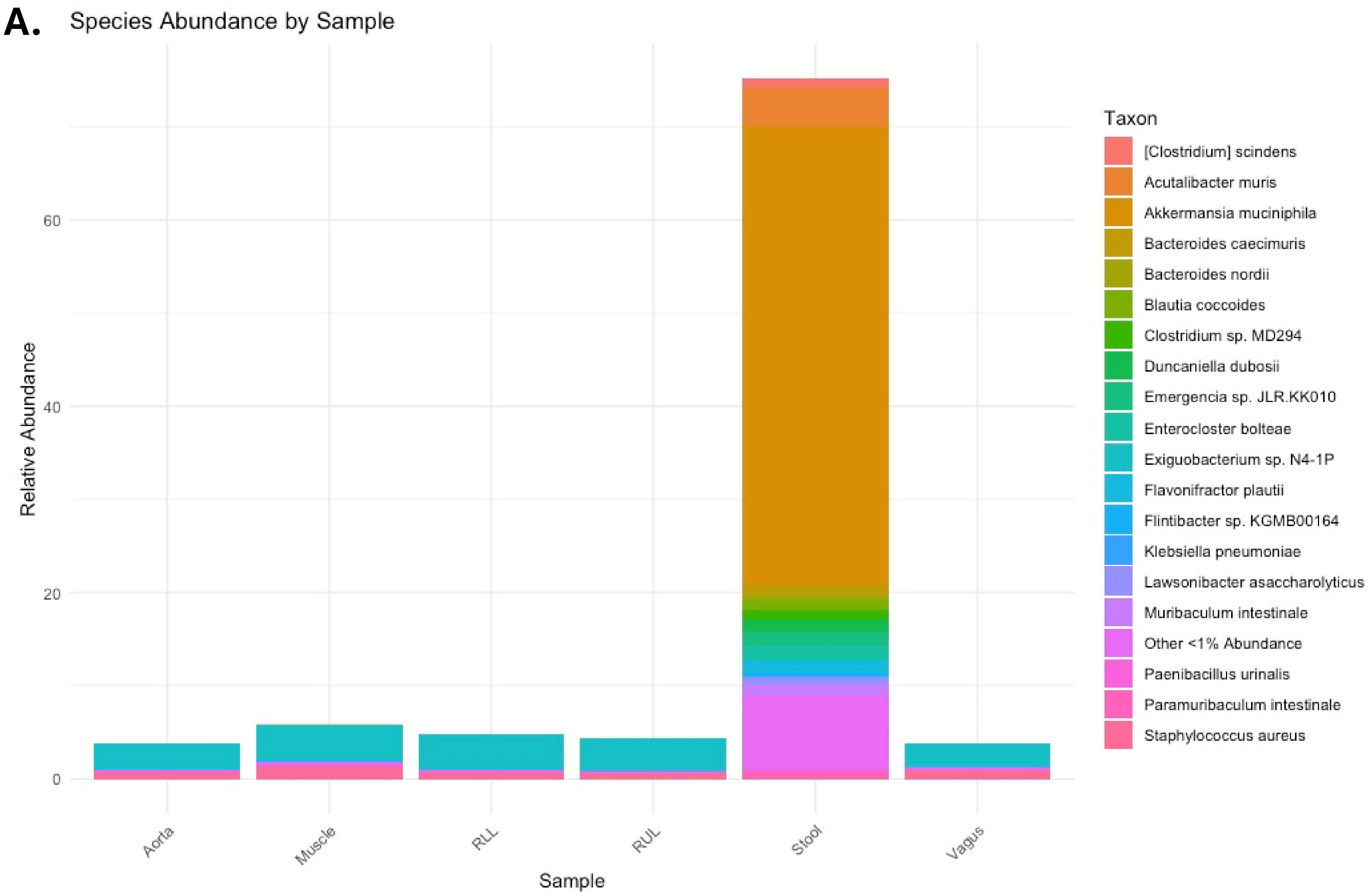

### Supplemental Figure 4

B.

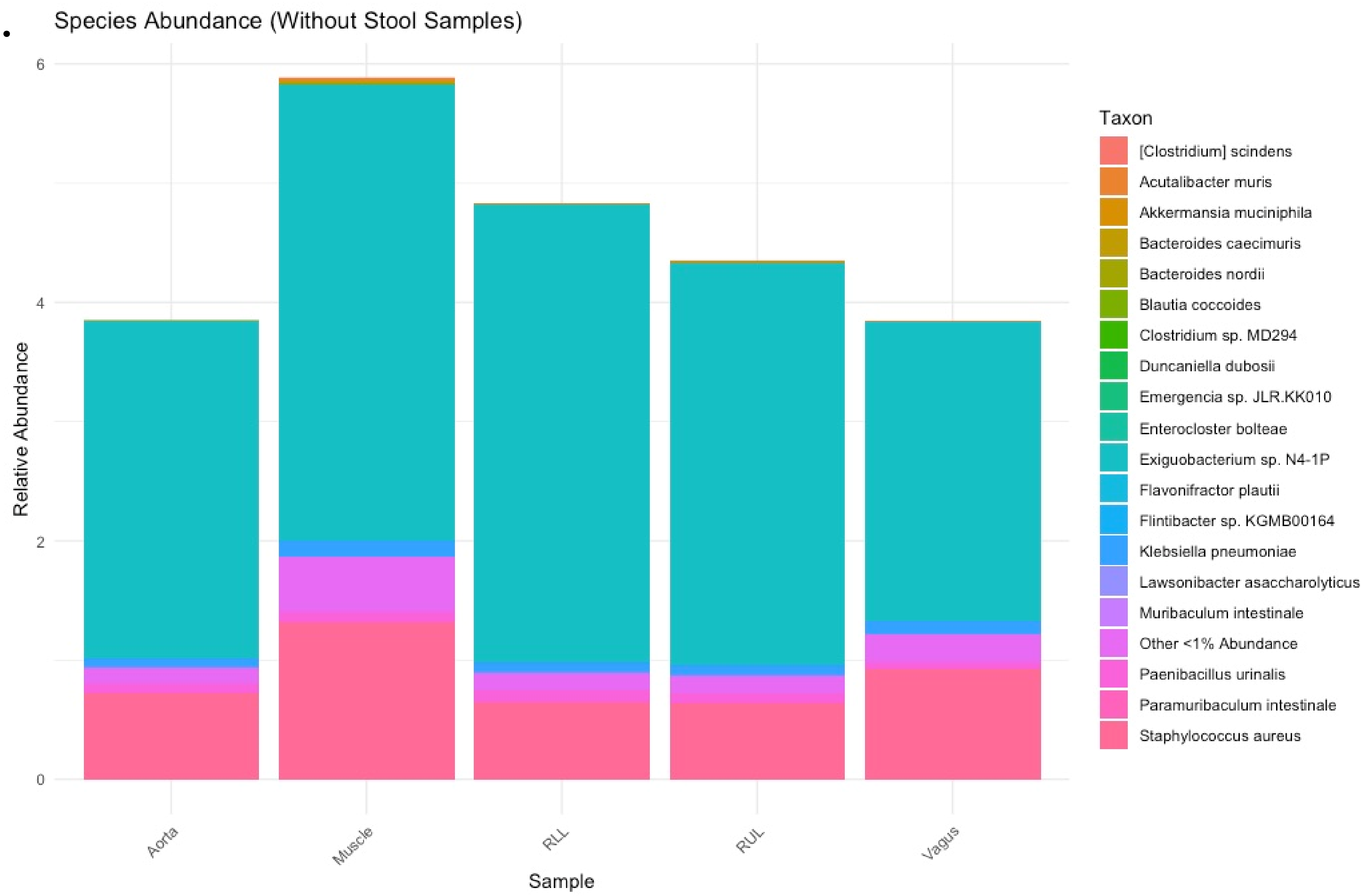

**Supplemental Table 1—CFU quantification of GFP-*E. coli* following gavage in germ free mice**

| Supplemental Table 1 (GF) | |
|---|---|
| **Organ** | **Mean Quantitative CFU Per Organ** |
| Stomach | 3.04E+07 |
| Jejunum | 3.02E+07 |
| Left Lung | 2.50E+06 |
| Right Lower Lung | 1.09E+06 |
| Right Upper Lung | 1.05E+06 |
| Right Middle Lung | 7.58E+05 |
| Esophagus | 6.48E+04 |
| Trachea | 5.68E+04 |
| AVN | 3.73E+04 |
| Heart | 2.22E+04 |
| CVN | 9.36E+03 |
| Stool | 8.50E+03 |
| Aorta | 2.55E+03 |
| Muscle | 6.88E+02 |
| Blood | 0.00E+00 |

**Supplemental Table 2—CFU quantification of GFP-*E. coli* following gavage in SPF mice**

| **Supplemental Table 2 (WT)** | |
|---|---|
| **Organ** | **Mean Qualtitative CFU Per Organ** |
| Jejunum | 4.56E+07 |
| Stomach | 4.01E+07 |
| Right Upper Lung | 6.48E+05 |
| Trachea | 5.52E+05 |
| Right Middle Lung | 5.50E+05 |
| Left Lung | 4.93E+05 |
| Esophagus | 4.26E+05 |
| Right Lower Lung | 3.15E+05 |
| Stool | 1.11E+05 |
| Heart | 8.18E+03 |
| AVN | 7.24E+03 |
| Aorta | 5.08E+03 |
| CVN | 5.43E+02 |
| Muscle | 9.72E+01 |
| Blood | 0.00E+00 |
